# Molecular classification of the placebo effect in nausea

**DOI:** 10.1101/2020.02.19.955740

**Authors:** Karin Meissner, Dominik Lutter, Christine von Toerne, Anja Haile, Stephen C. Woods, Verena Hoffmann, Uli Ohmayer, Stefanie M. Hauck, Matthias Tschöp

## Abstract

Numerous studies have shown that the mere expectation improvement can alleviate symptoms in various conditions. These ‘placebo effects’ often include reliable changes in central and peripheral organ systems. Here, we tested for the first time whether placebo effects can be monitored and predicted by plasma proteins. In a randomized controlled design, 90 healthy participants were exposed to a 20-min vection stimulus on two separate days and were randomly allocated to placebo treatment or no treatment on the second day. Significant placebo effects on nausea, motion sickness, and gastric activity could be verified. Using state-of-the-art proteomics, 74 differentially regulated proteins were identified in placebo-treated participants as compared to no-treatment controls. Gene ontology (GO) enrichment analyses of these proteins revealed acute-phase proteins as well as microinflammatory proteins to be reliable plasma correlates of the placebo effect. Regression analyses showed that day-adjusted scores of nausea indices in the placebo group were predictable by the identified GO protein signatures. We next identified specific plasma proteins, for which a significant amount of variance could be explained by the experimental factors ‘sex’, ‘group’, ‘nausea’, or their interactions. GO enrichment analyses of these proteins identified ‘grooming behavior’ as a prominent hit, based on ‘neurexin-1’ (NRXN1) and ‘contactin-associated protein-like 4’ (CNTNAP4). Finally, Receiver Operator Characteristics (ROC) allowed to identify specific plasma proteins differentiating placebo responders from non-responders. These comprised immunoglobulins (IGHM, IGKV1D-16, IGHV3-23, IGHG1) and MASP2, related to regulation of complement activation, as well as proteins involved in oxidation reduction processes (QSOX1, CP TXN). This proof-of-concept study indicates that plasma proteomics are a promising tool to identify molecular correlates and predictors of the placebo effect in humans.

## Introduction

The neurobiological mechanisms underlying placebo effects have become a research topic of increasing academic interest and intense study over the last decade. Approaches toward identification of exact mechanistic underpinnings frequently focus on changes in brain activity and brain connectivity to the release of neurotransmitters, including endogenous opioids, endocannabinoids, and dopamine^1, 2^. For the field of clinical research, a perhaps even more profitable approach would be the discovery of circulating biomarkers, which would provide the potential to predict placebo responders and monitor placebo effects in clinical trials without having to include cumbersome, expensive and invasive placebo control groups. However, to date, few studies have sought accessible markers for the placebo effect in blood, and they typically followed a narrow candidate-based approach^3^. Recent advances in high-throughput proteomics in combination with next-generation bioinformatics have now enabled novel ways to identify the molecular fingerprint of placebo effects in human plasma.

The purpose of this randomized controlled study was to determine whether the placebo effect can be tracked and predicted by proteins in peripheral blood. Our goal was to develop a method for the reliable and artifact-free verification of placebo effects on clinical endpoints reflecting broadly relevant disease symptoms and chose a well-established model of the placebo effect related to nausea^4, 5, 6^. In a randomized and carefully controlled design, 90 healthy participants were exposed for 20 minutes to a virtual vection drum on two separate days. On the second day after the baseline measurement, participants were randomly assigned to either placebo intervention (sham-TENS stimulation of a dummy acupuncture point) or to no treatment. Plasma samples for proteomics assessments were collected during baseline and at the end of the vection stimulus on both days following precise procedures for sample collection and handling.

## Methods

### Participants

Healthy right-handed volunteers between 18 and 50 years and of normal body weight and normal or corrected-to-normal vision and hearing were included. Exclusion criteria comprised metal implants or implanted device, the presence of acute or chronic disease, and regular intake of drugs (except for hormonal contraceptives, thyroid medications, and allergy medications). Furthermore, participants were excluded when they presented with anxiety and depression scores above the clinically relevant cut-off score according to the Hospital Anxiety and Depression Scale (HADS)^7^, when they scored lower than 80 in the Motion Sickness Susceptibility Questionnaire (MSSQ)^8^, or when they developed less than moderate nausea (<5 on 11-point NRS) in the pre-test session, during which participants were exposed to the nauseating vection stimulus for ≤ 20 minutes.

### Study design

All participants underwent a baseline session (Day 1) and a testing session (Day 2) on two separate days at least 24 hours apart. On Day 2 after the 20-min resting period, 60°participants (30 males and 30 females) were randomly assigned to placebo treatment, 30°participants (15 males, 15 females) to no treatment (Fig. 1a & 1b), and 10 participants to active treatment. The active treatment group (data not analyzed) was included to allow for the blinded administration of the placebo intervention, a common approach in placebo studies^9^. The no treatment group served to control the placebo effect for naturally occurring changes from Day 1 to Day 2. The study protocol was approved by the ethical committee of the Medical Faculty at Ludwig-Maximilians-University Munich (no. 402-13). All participants provided written informed consent. The study was registered retrospectively at the German Clinical Trials Register (no. DRKS00015192).

**Figure 1.**
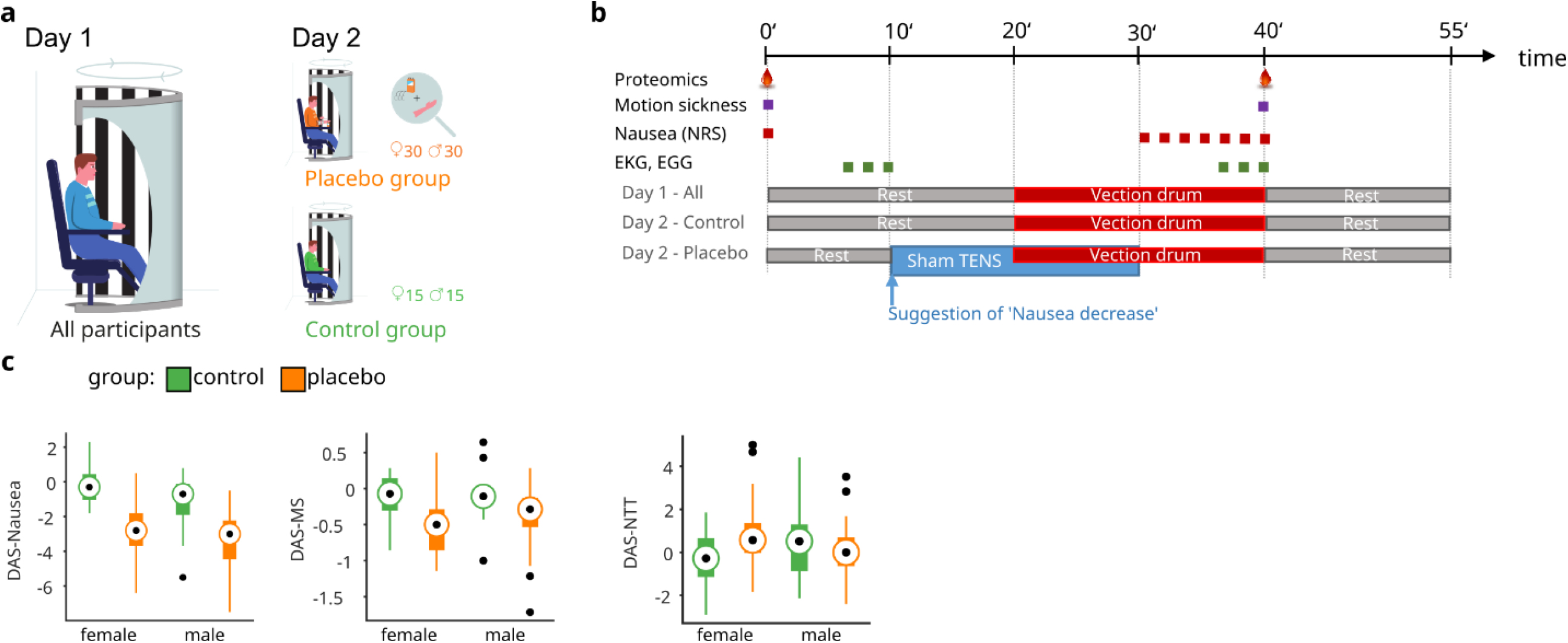
(**a)** Experimental design. (**b)** Experimental procedure. (**c)** Effect of placebo intervention on nausea. Boxplots depict day-adjusted scores (DAS) for the three nausea measures: Nausea (DAS-Nausea), motion sickness (DAS-MS), normo-to-tachy ratio in the electrogastrogram (DAS-NTT). DAS-Nausea: *F*_group_(1,86) = 44.83, P < 0.001; DAS-MS:_group_(1,84) = 14.93, P < 0.001; DAS-NTT: *F*_int_(1,83) = 4.16, P = 0.044 (female, *F*_group_(1,40) = 4.10, P = 0.049; male, *F*_group_(1,43) = 0.83, P = 0.366); See Supplemental Table 2 for further details. On each box, the central dot indicates the median, and the bottom and top edges of the box indicate the 25th and 75th percentiles, respectively. The whiskers extend to the most extreme data points not considered outliers, and the outliers are plotted as black dots.

Each participant was tested after a fasting period of at least 3 h on two separate days at least 24 h apart at the same daytime between 14.00 and 19.00 h. On Day 1, the procedure started with a 20-min resting period and then the vection stimulus was turned on for 20 min (Fig. 1b). On Day 2, after a 10-min baseline assessment, participants were randomly allocated to either placebo treatment, or active treatment, or no treatment. For subjects in the active and placebo treatment groups, a standardized expectancy manipulation procedure was performed, the electrodes were attached and the placebo or active TENS treatment was turned on for 20 minutes (Fig. 1b). On both days, heart rate, gastric myoelectrical activity, respiration frequency, the ECG, and a 32-lead electroencephalogram (EEG) were continuously recorded, and subjects rated the intensity of perceived nausea as well as other symptoms of motion sickness once each minute (Fig. 1b). For security reasons, the vection stimulus was stopped if nausea ratings indicated severe nausea (ratings of 9 or 10 on the NRS).

### Interventions

Placebo and active TENS interventions were implemented by means of a programmable TENS device (Digital EMS/TENS unit SEM 42, Sanitas, Uttenweiler, Germany). For the placebo treatment the electrodes were attached just proximal and distal to a non-acupuncture point at the ulnar side of the forearm generally accepted to represent a dummy point in the context of acupuncture research^10^. Two types of placebo stimulation were applied: 30 participants (15 males, 15 females) received subtle stimulation at a very low intensity by turning on the massage program of the TENS device (‘enhanced placebo’), while 30 participants (15 males, 15 females) received no electric stimulation at all (‘simple placebo’). Since the placebo effect did not differ between the two placebo groups^6^, the two placebo groups were merged. For the active intervention, the same device was used, but the electrodes were placed around ‘PC6’, a validated acupuncture point for the treatment of nausea^11, 12^, and the TENS program was turned on for 20 minutes.

### Randomization and blinding

Computer-assisted randomization was performed by a person not involved in the experiments, who prepared sequentially numbered, sealed and opaque randomization envelopes. The interventions were performed in a single-blind design. Participants in the no-treatment control group were necessarily unblinded.

### Nausea induction

Nausea was induced by standardized visual presentation of alternating black and white stripes with left-to-right circular motion at 60 degree/sec. This left-to-right horizontal translation induces a circular vection sensation wherein subjects experience a false sensation of translating to the left^13, 14^. The nauseating stimulus was projected to a semi-cylindrical and semi-transparent screen placed around the volunteer at a distance of 30 cm to the eyes. Such stimulation simulates visual input provided by a rotating optokinetic drum, commonly used to induce vection (illusory self-motion) and thereby nausea^15, 16^.

### Proteomic analyses

Plasma samples were proteolysed using PreOmics’ iST Kit (PreOmics GmbH, Martinsried, Germany) according to manufacturers’ specifications. After drying, the peptides were resuspended in 2% ACN and 0.5% TFA acid. The HRM Calibration Kit (Biognosys, Schlieren, Switzerland) was added to all of the samples according to manufacturer’s instructions.

Mass spectrometry data were acquired in data-independent acquisition (DIA) mode on a Q Exactive (QE) high field (HF) mass spectrometer (Thermo Fisher Scientific Inc.). Per measurement 0.5 μg of peptides were automatically loaded to the online coupled RSLC (Ultimate 3000, Thermo Fisher Scientific Inc.) HPLC system. A nano trap column was used (300 μm inner diameter (ID) × 5 mm, packed with Acclaim PepMap100 C18, 5 μm, 100 Å; LC Packings, Sunnyvale, CA) before separation by reversed-phase chromatography (Acquity UPLC M-Class HSS T3 Column 75µm ID x 250mm, 1.8µm; Waters, Eschborn, Germany) at 40°C. Peptides were eluted from the column at 250 nl/min using increasing acetonitrile (ACN) concentration (in 0.1% formic acid) from 3% to 40 % over a 45-min gradient.

The HRM DIA method consisted of a survey scan from 300 to 1500 m/z at 120,000 resolution and an automatic gain control (AGC) target of 3e6 or 120 msec maximum injection time. Fragmentation was performed via high-energy collisional dissociation (HCD) with a target value of 3e6 ions determined with predictive AGC. Precursor peptides were isolated with 17 variable windows spanning from 300 to 1500 m/z at 30,000 resolution with an AGC target of 3e6 and automatic injection time. The normalized collision energy was 28 and the spectra were recorded in profile type.

Data analysis of DIA files requires comparison of mass spectra against a tailored spectral library built of preceding data dependent mass spectrometry measurements. We searched our DIA files against an in-house library generated from selected mass spectrometry data encompassing 57 files of plasma and serum preparations, spiked with the HRM Calibration Kit (Biognosys). Data dependent files were analyzed using Proteome Discoverer (Version 2.1, ThermoFisher Scientific). Embedded search engine nodes included Mascot (Version 2.5.1, Matrix Science, London, UK), Byonic (Version 2.0, Proteinmetrics, San Carlos, CA), Sequest HT, and MSAmanda. Peptide FDRs for all search engines were calculated using percolator node, and the resulting identifications were filtered to satisfy the 1% peptide level FDR (with the exception of Byonic) and combined in a multi-consensus result file maintaining the 1% FDR threshold. The peptide spectral library was generated in Spectronaut (Version 9, Biognosys) with default settings using the Proteome Discoverer combined result file. Spectronaut was equipped with the Swissprot human database (Release 2016.02, 20165 sequences, www.uniprot.org) with a few spiked proteins (e.g., Biognosys iRT peptide sequences). The final spectral library generated in Spectronaut contained 1811 protein groups and 26,805 peptide precursors.

The DIA mass spectrometry data were analyzed using the Spectronaut 9 software applying default settings with the following exceptions: quantification was limited to proteotypic peptides, data filtering was set to Qvalue sparse for the peptide-based analysis. The Qvalue sparse setting includes all observations that pass the Qvalue at least once and it generates a matrix with a minimum of missing values. A peptide dataset used for ANCOVA was generated by removing peptide with more than 10% missing values. Then data were normalized by setting the median to one, and intensities were log transformed (logN) for further analyses. The final peptide dataset consisted of 16,180 entries.

For the protein-based analysis, unique peptides were merged to proteins. To ensure best data quality in a most complete protein matrix, data were filled up in two steps. First, data filtering was set to Qvalue. The Qvalue setting considers only individual observations that pass the Qvalue threshold and generates a matrix containing missing values. Proteins with >95% missing values were deleted. Instead of imputing the remaining missing data, we used the values generated from the Qvalue sparse setting, representing real mass traces at the respective retention times. Data were finally normalized setting the median to one, intensities were log transformed (logN) for further analyses. The dataset finally consisted of 718 unique proteins. We identified 2°samples as haemolytic, which were removed from both datasets for all analyses. Fold changes (PFC) were calculated for both days as log2 fold between measurement 2 and 1.

### Behavioral and physiological data

Perceived nausea intensities were rated at baseline and every minute during the nausea period on 11-point NRSs, with ‘0” indicating ‘no nausea” and ‘10” indicating ‘maximal tolerable nausea.”. Symptoms of motion sickness were assessed by using the ‘Subjective Symptoms of Motion Sickness’ (SSMS) questionnaire adapted from^8^, with scores of 0 to 3 assigned to responses of none, slight, moderate, and severe for symptoms of dizziness, headache, nausea/urge to vomit, tiredness, sweating, and stomach awareness, respectively.

The EGG signal, respiratory activity (to control the EGG for respiratory artifacts), and the ECG signal (results not reported here) were recorded using a BIOPAC MP 150 device (BIOPAC Systems Inc., Goleta, CA, USA) and AcqKnowledge 4.1 software for data acquisition. The EGG signal was recorded by using two Ag/AgCl electrodes (Cleartrace, Conmed, Utica, NY, USA) placed at standard positions on the skin above the abdomen^17^. The skin was cleaned with sandy skin-prep jelly to reduce skin impedance prior to electrode placement (Nuprep, Weaver & Co., Aurora, CO, USA). The electrodes were connected to the BIOPAC amplifier module EGG100C, the signal was digitized at a rate of 15.625 samples per second and filtered using an analog bandpass filter consisting of a 1 Hz first-order low-pass filter and a 5 MHz third-order high-pass filter. Spectral analysis was performed on the last 300 sec of the baseline and nausea periods on each testing day, respectively (Fig. 1b). Prior to fast Fourier transform (FFT), each data epoch was linearly detrended and its ends were tapered to zero using a Hamming window. Spectral power within the normogastric bandwidth (2.5 to 3 cycles per min) and the tachygastric bandwidth (3.75 to 9.75 cycles per min) were determined for three overlapping epochs (Minutes 1 to 3, 2 to 4, 3 to 5)^18^. Finally, the average ‘normo-to-tachy ratio’ (NTT) was computed as the mean ratio of normogastric to tachygastric spectral power in the three 1-min epochs. NTT is known to decrease during visually-induced nausea, indicating enhanced tachygastric myoelectrical activity and/or reduced normogastric myoelectrical activity^19, 20, 21^. NTT data were logarithmized before statistical analysis to obtain normal distributions.

The EOG was recorded to control for participant’s eye movements during baseline and vection stimulation in order to assure that they followed the standardized instructions, namely to look straight ahead during baseline and to naturally follow the left-to-right horizontal translation of black and white stripes without moving the head during exposure to the vection stimulus, respectively. Horizontal and vertical electrooculography (EOG) was assessed using the ActiveTwo system (BioSemi, Amsterdam, NL).

### Statistical analysis

For all statistical tests, a significance level of *P* ≤ .05 (two-tailed) was assumed. For each behavioral and physiological variable (nausea, MS, NTT), day-adjusted scores (DAS) were computed prior to statistical analyses. DAS were calculated as: DAS = (m_22_ – m_21_) – (m_12_ – m_11_), where m_22_ is measurement 2 on Day 2, m_21_ is measurement 1 on Day 2 and m_12_ and m_11_ are second and first measurement on Day 1, respectively. DAS for each nausea, MS, and NTT were subjected to separate analyses of variance (ANOVAs), with ‘group’ (placebo group, control group) and ‘sex’ (male, female) included as between-subject factors. Assumptions of normality were verified for all outcomes before statistical analysis.

To adjust for individual differences in protein fold changes during the baseline experiment on Day 1, we applied an analysis of covariance (ANCOVA) to estimate an unbiased difference in placebo-induced protein abundances on Day 2. To further increase statistical power, ANCOVA was performed on the log ratios (measurement 2 vs measurement 1) of precursors instead of protein data. Thus, for large individual proteins we could increase the sample size up to 1000 fold (supplemental Table 1).

**Table 1.** Sample characteristics at baseline.

|  | No treatment<br>(n=30) | Placebo treatment<br>(n=60) | *P* * |
| --- | --- | --- | --- |
| Sex, *m/f* | 15/15 | 30/30 | 1 |
| Age, *mean (s.d.)* | 23.5 (2.7) | 23.5 (3.4) | .49 |
| Education (≥high school degree), *n (%)* | 27 (90) | 59 (98%) | .58 |
| Nonsmoker, *n (%)* | 26 (87%) | 53 (90%) | .73 |
| Body Mass Index, *mean (s.d.)* | 22.3 (2.7) | 21.6 (2.1) | .18 |
| MSSQ, *mean (s.d.)* | 130.3 (38.4) | 137.1 (39.9) | .44 |
| HADS-anxiety, *mean (s.d.)* | 4.0 (2.6) | 4.0 (2.2) | .95 |
| HADS-depression, *mean (s.d.)* | 1.4 (1.6) | 1.8 (1.6) | .35 |
| STAI-state anxiety, *mean v* | 34.7 (4.9) | 35.8 (8.3) | .51 |
| STAI-trait anxiety, *mean v* | 38.8 (6.4) | 37.8 (6.5) | .51 |
| PSQ-Stress, *mean v* | 31.1 (14.3) | 30.0 (16.1) | .72 |

Abbreviations: MSSQ, Motion Sickness Questionnaire; HADS, Hospital Anxiety and Depression Scale; STAI, State-Trait Anxiety Inventory; PSQ, Perceived Stress Questionnaire. \*Results of $\chi^2$ tests, Kruskal-Wallis tests, or univariate ANOVA, as appropriate.

The protein interaction network was created using StringDB Version 1019 using default settings (median confidence). Edges refer to the interaction sources text-mining, experiments, databases, co-expression, neighborhood and co-occurrence. GO Enrichment analyses were done using GO annotation files from http://www.geneontology.org/, releases/2017-03-11. Significantly enriched GO terms were estimated using a hypergeometric distribution test with the full Proteomics Spectronaut 9 Database (1470 unique protein IDs) as background. GO terms are highly redundant. Fully redundant terms were removed from the list. To form representative functional clusters, similar terms were combined using the Jaccard index J_ij=(g_i∩g_j)⁄(g_i∪g_j), where gi and gj are the gene-products assigned to significant enriched pairs of GO-terms i and j, respectively. GO-terms with an index Jij > 0.4 were grouped.

For each GO term enriched for the ANCOVA-identified proteins we selected all related proteins that were also significantly regulated and generated two linear regression models to predict the three DAS (Nausea, NTT, MS) for the control and placebo groups. A ‘bisquare’ weight function was used to remove outliers from the model. To estimate the variance composition of the protein changes on Day 2 dependent on the three DASs, we used a linear regression model: *y*_*p,NS*_= *μ* + *β*_1_ ∗ *DAS* + *β*_2_ ∗ *grp* + *β*_3_ ∗ *sex* + *β*_4_ ∗ *DAS* ∗ *grp* + *β*_5_ ∗ *DAS* ∗ *sex* + *β*_6_ ∗ *grp* ∗ *sex* + *β*_7_ ∗ *DAS* ∗ *grp* ∗ *sex*, with DASs as the predictor variables and group (grp) and sex as categorical co-variables. We used ‘bisquare’ weight function to detect and remove outliers from the model. The proportion of total variance explained by the regression model was estimated by comparing the regression sum of squares to total sum of squares. Variance composition was computed separately for each imputed dataset. Results were summarized by taking the medians for each protein.

Prediction of placebo responders was done based on protein baseline data on Day 2. We performed one-way ANOVA on protein level to preselect protein differentially expressed between ‘placebo responders’ (i.e., showing ≥50% reduction in DAS-nausea and DAS-MS, respectively) and all other participants (‘non-responders”) in the placebo group. Additionally, we performed sequential feature selection (SVS) to identify proteins with potential to predict good responders. We finally used a linear support vector (SVM) machine to generate two predictive models for both Nausea scores, one including ANOVA and SVS predictor proteins (‘ANOVA plus model’) and a model based on randomly selected proteins (‘RANDOM model’). Median ROC curves and mean AUC estimation were done using k-fold cross validation with k = 10, with 10 independent permutations. Statistics were done using MATLAB R2018b.

## Results

### Participant characteristics

Hundred participants were randomized (60 placebo TENS, 30 no treatment, 10 active TENS). Analyses were based on the data from 90 participants assigned to placebo treatment or no treatment (Fig. 1a). Groups were comparable at baseline with regard to demographic and clinical characteristics (Table 1).

### Nausea and motion sickness

As expected, the placebo-exposed individuals developed fewer symptoms of nausea and motion sickness on Day 2 (Fig. 1c, supplemental Table 1). Consistent with this observation, analyses of variance (ANOVA) revealed a significantly larger reduction in day-adjusted nausea scores (DAS-nausea) from Day 1 to Day 2 in the placebo group as compared to the control group (*F*_group_(1,86) = 44.83, *P* < 0.001), with no difference between sexes (*F*_int_(1,86) = 1.01, *P* = 0.32). After removal of one outlier (Fig. 1c) the main effect of ‘group’ remained significant (*P* < 0.001). Thirty-nine out of 60 patients (65%) in the placebo group and 5 out of 30 (17%) in the control group (*χ*^2^ = 18.69, *P* < 001) showed a reduction of at least 50% in DAS-nausea.

Similarly, the ANOVA for day-adjusted motion sickness scores (DAS-MS) indicated relief from MS in the placebo group as compared to the control group (*F*_group_(1,84) = 14.93, P < 0.001), with no difference between sexes (*F*_int_(1,84) = 0.47, *P* = 0.49). After removal of six outliers (Fig. 1c) the main effect of ‘group’ remained significant (*P* < 0.001). Thirty out of 58 patients (52%) in the placebo group and 3 out of 29 (10%) in the control group showed a reduction in DAS-MS of at least 50%, the difference between groups was significant (*χ*^2^ = 14.06, *P* < 0.001).

The ANOVA for the day-adjusted changes in the normo-to-tachy ratio in the electrogastrogram (DAS-NTT), revealed a significant interaction between ‘group’ and ‘sex’ (*F*_int_(1,83) = 4.16, *P* < 0.05). Contrast analyses indicated a significant difference among treatment groups for female participants (*F*_group_(1,40) = 4.10, *P* < 0.05), but not for males (*F*_group_(1,43) = 0.83, *P* = 0.40; Fig. 2a). After removal of four outliers (Fig. 1c) the interaction between ‘group’ and ‘sex’ remained significant (*P* < 0.05).

**Figure 2.**
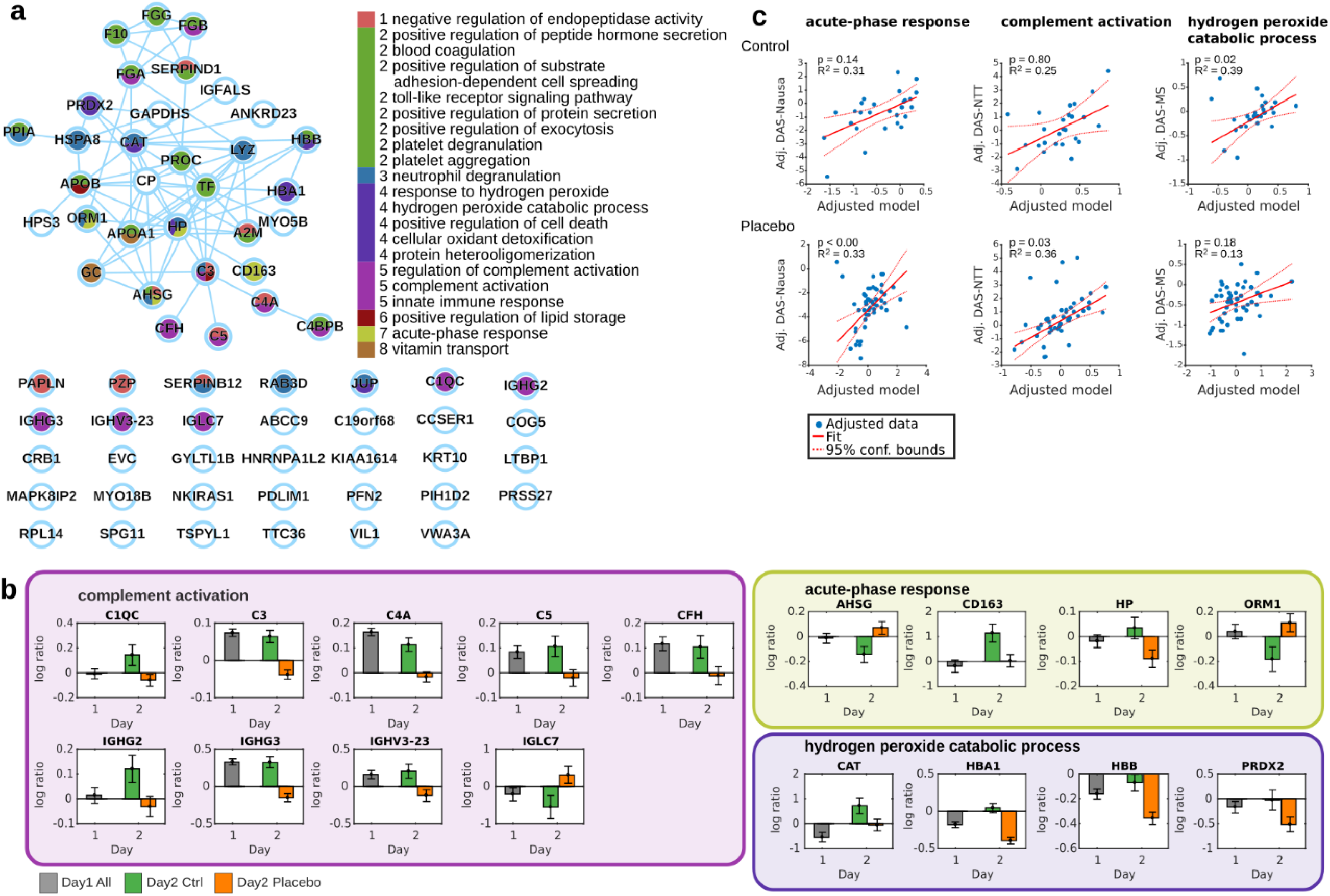
**(a)** String DB network of placebo-related proteins. Nodes are colored according to GeneOntology term group association. Numbers refer to functional clusters. GO functional clusters were created based on shared members (proteins). Edges refer to String DB interactions. (see methods). **(b)** Expression patterns for selected GO term proteins. Barplots depict average log expression (measurement 2 vs measurement 1) of all protein-associated peptide precursors. Barplot colors refer to day and group. Error bars denote standard error of the mean. Boxes around barplots indicate GO term membership, and the color of the boxes refer to GO functional cluster associated in panel a. **c)** The fitted GO term-based linear multiple regression models to predict DAS. DAS-Nausea predicted by acute-phase response proteins in control (top) and placebo group (bottom); complement activation proteins predict DAS-NTT in control and placebo; hydrogen peroxide catabolic process proteins predict DAS-MS in control and placebo; Model P-values (FDR-corrected) and model R-squared are specified in the plots. Blue dots are model outputs for each data point. The linear fit and 95% confident bands are denoted by solid and dashed red lines.

### Proteins in peripheral plasma

A mass spectrometry-based proteomics approach on peripheral plasma identified 711 proteins represented by 3224 peptides and 14588 peptide-precursors in at least 25 percent of participants, with many proteins showing a marked sex difference in abundance, but no such influence of age, group, day, or time point of measurement (supplemental Fig.).

### Placebo proteome

To identify specific placebo-associated protein changes, analyses of covariance (ANCOVA) were performed for vection-induced fold changes of protein precursors/peptides (PFC) on Day 2 including the factors ‘group’ and ‘sex’ as between-subject factors and PFCs on Day 1 as covariates. A significant main effect of ‘group’ was found for 74 proteins (Fig. 2b; supplemental Table 3), indicating differential regulation of these proteins on Day 2 in the placebo group as compared to the control group. In more detail, and relative to the controls, 34°proteins were more abundant following placebo treatment and 30 were less abundant. Mapping these proteins to the StringDB^22^ revealed a functional network of 33 proteins as well as 31 unconnected proteins (the remaining 10 proteins could not be mapped uniquely; Fig. 2a).

### Gene ontology (GO) enrichment analyses of the placebo proteome

We next performed gene ontology (GO) enrichment analyses to identify the predominant biological processes, in which these 74 placebo-related proteins are involved functionally. Thirty-three enriched GO terms could be identified, from which 21 non-redundant terms could be grouped into 8 functional clusters of 37 proteins (FDR-corrected *P* < 0.05; supplemental Table 4). The most striking protein pattern was detected for the GO term ‘complement activation’, with 8 out of 9 proteins having an equal abundance pattern, namely decrease in the placebo group as compared to the control group (Fig. 2b). Involved proteins were complement C3, C4a and C5, complement factor H (CFH), complement C1q subcomponent subunit C (C1QC), immunoglobulin heavy variable 3-23 (IGHV3_23), and immunoglobulin heavy constant gamma 2 and 3 (IGHG2, IGHG3).

### Prediction of placebo effects by GO terms

To evaluate the predictive value of GO term related proteins for nausea indices, we performed regression analyses for each DAS nausea score in the placebo and control groups, respectively. In the placebo group, DAS-Nausea was best predicted by PFC of proteins involved in the acute phase-response, namely haptoglobin precursor (HP), alpha-1-acid glycoprotein 1 (ORM1), alpha-2-HS-glycoprotein (AHSG), and ‘scavenger receptor cysteine-rich type 1 protein M130’ (CD163). Furthermore, DAS-NTT was predicted by proteins related to the GO terms ‘blood coagulation’ and ‘complement activation’ (Fig. 2c; supplemental Table 5). Key proteins for the association with blood coagulation were fibrinogen alpha and beta (FBA and FBB; supplemental Table 6). In the control group, DAS-MS was predicted by the GO terms ‘hydrogen peroxide catabolic process’, ‘positive regulation of substrate adhesion-dependent cell spreading’, and ‘protein heterooligomerization’ (Fig. 2c; supplemental Table 6). Key proteins for these GO terms were apolipoprotein A-I (APOA1), hemoglobin subunit beta (HBB), and hemoglobin subunit alpha (HBA1).

### Dissection of protein variance by experimental factors

In a second approach, we dissected the variance of PFC on Day 2 and determined to what extent it was determined by the experimental factors ‘group’, ‘sex’, ‘DAS-Nausea’ (or ‘DAS-MS’, ‘DAS-NTT’), or by the interaction terms. Proteins for which a significant amount of variance could be explained are summarized in Fig. 3a (see also supplemental Tables 6, 7, 8). GO enrichment analyses of these proteins were performed separately for each type of nausea score (supplemental Tables 9, 10, and 11, respectively). One of the most significant hits was the GO term ‘grooming behavior’ in models including ‘DAS-NTT’ (Fig. 3b, supplemental Table 11) with the key proteins ‘neurexin-1’ (NRXN1) and ‘contactin-associated protein-like 4’ (CNTNAP4). A further key protein in models including ‘DAS-NTT’ was reelin (RELN) (supplemental Table 11).

**Figure 3.**
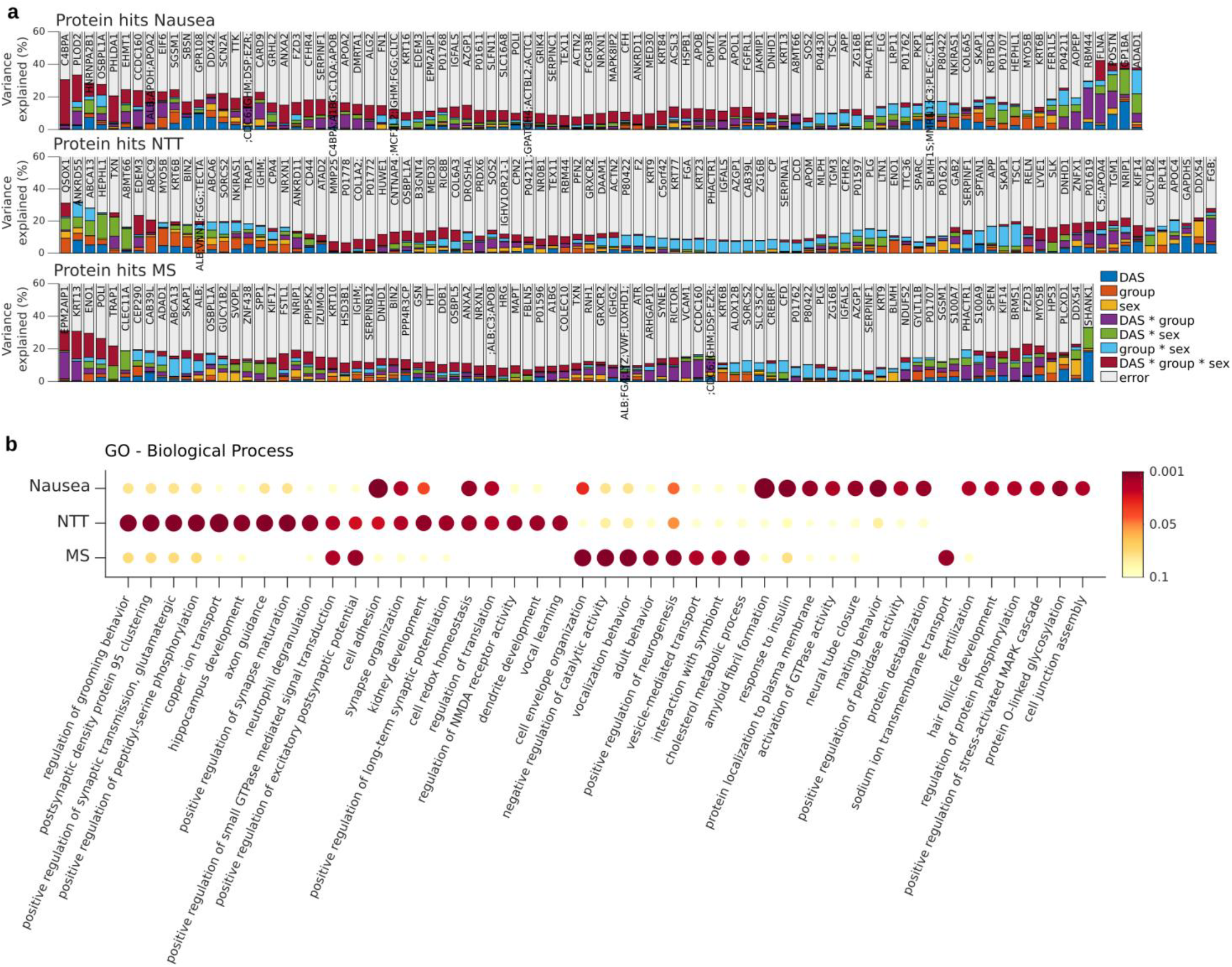
**(a)** Variance composition of proteins. The amount of variance explained was estimated using linear multiple regression models to predict protein fold change variance by different experimental factors (DAS, sex, group) and their interactions (predictor variables). Models were generated independently for each protein and type of nausea measure: Nausea, NTT and MS. Barplot histograms depict variance composition for all proteins significantly affected (P < 0.05) by at least one predictor variable. Multiple protein labels arise from non-unique mapped precursors. **(b)** GO enrichment for each group of significantly regulated proteins (P < 0.05). Dot color and size refer to FDR-corrected enrichment –log10 (p-value).

### Responder analysis

A one-way ANOVA on protein level was performed to preselect proteins at baseline of Day 2 that were expressed differentially between placebo responders and placebo non-responders for nausea and motion sickness, respectively. AUC estimators for the ANOVA plus model were 0.86 for nausea and 0.93 for motion sickness, respectively, compared to 0.6 ± 0.07 for the random model (Fig. 4a, 4b). Proteins differentiating between placebo responders and placebo-non-responders comprised immunoglobulins (IGHM, IGKV1D-16, IGHV3-23, IGHG1) and MASP2, which are related to regulation of complement activation, as well as proteins related to oxidation reduction processes (QSOX1, CP TXN). Also included was SLC9A3R1, a protein related to various biological processes, including dopamine receptor signaling (supplemental Tables 12, 13).

**Figure 4.**
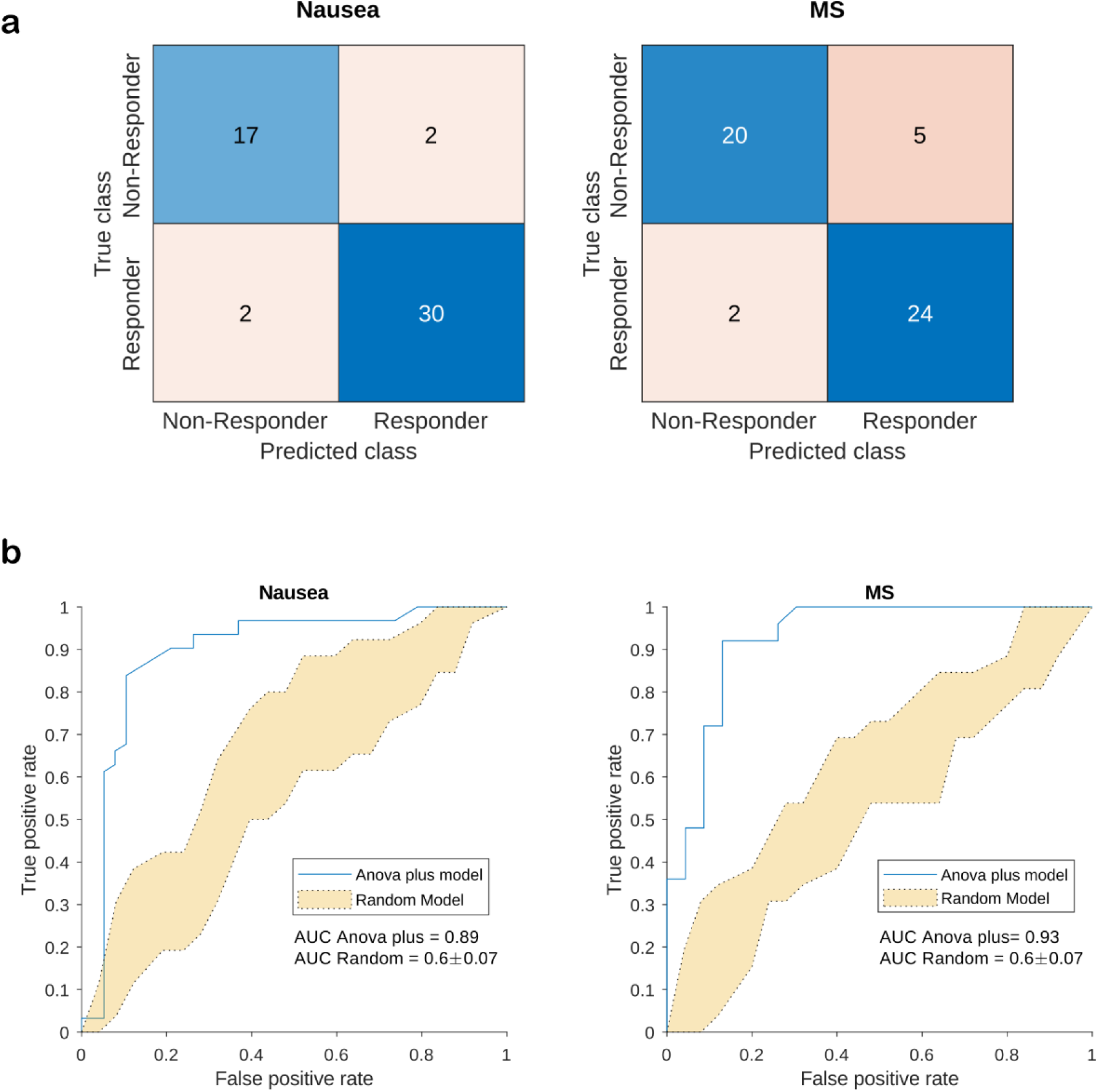
**(a)** Adapted confusion matrices for placebo responders vs. non-responders in the placebo group for nausea and motion sickness (MS). Placebo responders were defined as participants showing a reduction of at least 50% in DAS-nausea and DAS-MS, respectively. **(b)** Receiver operator characteristics (ROC) for the SVM models. The blue line refers to the ANOVA plus model. Yellow area refers to the range of all 10 RANDOM model permutations, AUC values in mean and standard deviation given.

## Discussion

Moving beyond recent studies, which used genomic techniques to uncover the mechanistic basis of placebo effects^23, 24^, we here discovered for the first time specific molecular signatures reflecting the placebo effect and its impact in acute nausea. By applying next generation computational and bioinformatics approaches, we identified distinct biological processes that were associated with the placebo effects. For example, the acute-phase proteins HP, ORM1, AHSG, and CD163 were reduced during placebo treatment and these changes were related to the decrease of nausea in the placebo group. The placebo-associated reduction of FBA and FBB, and the strikingly unique pattern regarding decrease of complement cascade proteins, fit well into this scheme because these indicators of microinflammation are known to increase in response to acute stress^25, 26, 27, 28, 29, 30, 31, 32, 33^. Indeed, an early study reported reduction of the acute-phase protein CRP (C-reactive protein) in placebo-treated surgery patients as compared to untreated controls^31^ and a recent hypothesis paper postulated that placebo effects are mediated by the suppression of the acute-phase response^29^. Our placebo intervention may thus have dampened microinflammatory processes related to acute nausea and motion sickness.

In a second approach, we aimed to predict the protein fold changes by changes in nausea and the experimental factors sex and group. Proteins most closely associated with the placebo effect in this model were led by neuropeptides that play a key role in social attachment and affiliation, including NRXN1, CNTNAP4, and RELN. NRXN1 and CNTNAP4 are both cell adhesion molecules involved in mirror neuron activity and empathic behavior ^34, 35, 36, 37, 38^, and RELN has been reported to functionally interact with oxytocin and both neuropeptides have been implicated in autism pathophysiology^39, 40^. These findings are consistent with earlier studies where administration of oxytocin and vasopressin, prior to eliciting a placebo effect, considerably increased the size of placebo effects in healthy volunteers^41, 42^. Most strikingly, one of our main hits in the GO enrichment analysis was ‘grooming behavior’. Grooming in various species has been postulated to constitute an important evolutionary trace of the placebo effect in humans^43, 44, 45^.

Finally, results indicate that plasma proteomics could be groundbreaking for the identification of biomarkers predicting placebo responders in clinical trials. ROC analyses revealed a protein pattern at baseline of Day 2 that allowed differentiating placebo responders from non-responders with surprisingly high accuracy (Fig. 4a, 4b). This set of proteins comprised immunoglobulins (IGHM, IGKV1D-16, IGHV3-23, IGHG1) and serum proteases (MASP2), both involved in the regulation of complement activation. Interestingly, changes of proteins related to this pathway were also significantly associated with the size of the gastric placebo effect (supplemental Table 6). Also included were proteins related to oxidative stress reduction (QSOX1, CP, TXN). Furthermore, SLC9A3R1 was part of the predictors, a protein involved also in dopamine receptor binding. The dopaminergic system gets activated before and during placebo interventions^46, 47, 48^. In addition, catechol-o-methyltransferase-gene *(COMT*) variants as well as personality traits related to the dopaminergic system explained a significant amount of variance of the placebo effect^49, 50^.

Some limitations of our study need to be acknowledged. First, certain circulating proteins may not have been identified in our proteome analysis, due to limitations in sensitivity. To increase reliability of results we did not focus on single protein hits but rather analyzed variations in GO pathways to detect changes in biologically connected patterns. This strategy successfully identified several biological processes, and importantly, these same processes had been shown to be relevant to placebo phenomena^29, 31, 50^. Secondly, we pursued an unbiased global proteomics approach to uncover a blood plasma protein signature imprinted by the placebo effect in humans and provided proof-of-principle evidence that circulating proteins predict and reflect the placebo effect in humans. However, we recognize that our results have to be regarded preliminary as long as validation studies are missing.

In sum, our results indicate that plasma proteomics is a timely and promising approach to quantify and predict the placebo effect in peripheral blood. Our novel discovery of a proteomic fingerprint of placebo effects in peripheral blood thus offers transformative potential not only for a better understanding of the molecular basis of the placebo effect in different conditions but also for advancing and simplifying certain categories of clinical research in the future. Placebo arms in clinical trials are scientifically necessary for sound academic research, but they can be ethically irresponsible when novel therapeutics, for example in oncology, offer unprecedented and game-changing benefits. One considerable advantage of precision biomarkers based on plasma proteins is that peripheral blood is easily accessible. Once successfully validated across diseases, the inclusion of placebo control groups in clinical trials may no longer be necessary.

## Author contributions

K.M., S.H., M.T. designed the study, K.M., A.H., V.H. performed and analyzed the behavioral experiment, C.T., U.O., and S.H. performed the proteomic analysis, K.M., D.L., C.T., U.O., S.W., S.H., M.T. analyzed and interpreted the data, K.M., D.L., S.H., M.T. wrote the manuscript.

## Acknowledgments

We would like to thank Franziska Stahlberg and Simone Aichner for their valuable support in conducting the nausea experiment. The study was supported by a grant from the German Research Foundation (ME3675/1-1).

