## Supplemental Figure & Tables 2-4, 6-13 for "Molecular classification of the placebo effect in nausea"

### Supplemental Material

#### Supplemental Figure

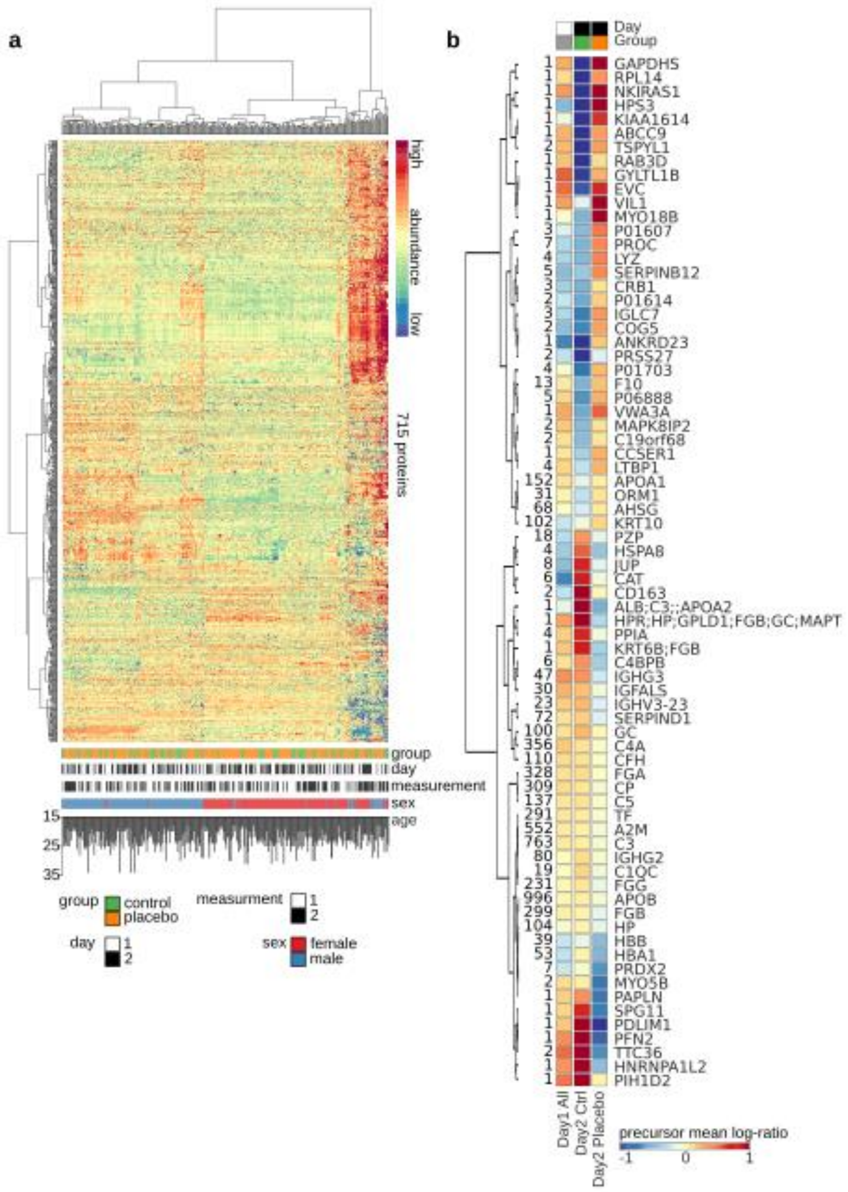

**Supplemental Figure. Identified plasma proteins and those significantly affected by the placebo treatment**. **(a)** The heatmap depicts the sample (column) vs protein (row) matrix with hierarchical clustering dendrograms. Abundances were row-wise z-score transformed. The sample features group, experiment day, time point of measurement, sex and age are color-/bar-coded at the bottom of the heatmap. **(b)** Heatmap of the 74 proteins identified as significantly affected ($P < 0.05$) by placebo treatment using ANCOVA, with fold changes on Day 1 included as covariates. Numbers on the left indicate numbers of peptide precursors associated with the protein. Color refers to average log-ratio (measurement 2 vs measurement 1) of all protein-associated peptide precursors. Columns are labeled with the same color-code as in A. Multiple protein labels arise from non-unique mapped precursors.

**Supplemental Table 2:** ANOVA results for day-adjusted scores (DAS) of nausea, motion sickness, and normo-to-tachy ratio (NTT).

| Measure | | Control group | | | Placebo group | | | 2-way ANOVA | | |
|---|---|---|---|---|---|---|---|---|---|---|
| | | *n* | *mean* | *s.d.* | *n* | *mean* | *s.d.* | *group* | *sex* | *group by sex* |
| DAS-Nausea (NRS 0-10) | | | | | | | | $F_{(1,86)}$ = 44.83, $P$ < 0.001 | $F_{(1,86)}$ = 3.05, $P$ = 0.084 | $F_{(1,86)}$ = 1.01, $P$ = 0.318 |
| | female | 15 | -0.32 | 1.33 | 30 | -3.10 | 1.63 | | | |
| | male | 15 | -1.31 | 1.64 | 30 | -3.37 | 1.71 | | | |
| | total | 30 | -0.82 | 1.55 | 60 | -3.23 | 1.66 | | | |
| DAS-MS | | | | | | | | $F_{(1,84)}$ = 14.93, $P$ < 0.001 | $F_{(1,84)}$ = 0.21, $P$ = 0.648 | $F_{(1,84)}$ = 0.47, $P$ = 0.493 |
| | female | 14 | -0.10 | 0.34 | 29 | -0.53 | 0.44 | | | |
| | male | 15 | -0.12 | 0.38 | 30 | -0.42 | 0.43 | | | |
| | total | 29 | -0.11 | 0.36 | 59 | -0.48 | 0.44 | | | |
| DAS-NTT (logratio) | | | | | | | | $F_{(1,83)}$ = 0.51, $P$ = 0.476 | $F_{(1,83)}$ = 0.0, $P$ = 0.996 | $F_{(1,83)}$ = 4.16, $P$ = 0.044 |
| | female | 13 | -0.18 | 1.41 | 29 | 0.82 | 1.51 | | | |
| | male | 15 | 0.56 | 1.69 | 30 | 0.08 | 1.65 | | | |
| | total | 28 | 0.22 | 1.58 | 59 | 0.44 | 1.61 | | | |

**Abbreviations:** DAS, day-adjusted score; NRS, numeric rating scale; bpm, beats per minute; MS, motion sickness; NTT, normo-to-tachy ratio

**Supplemental Table 3:** 74 proteins that were differentially regulated in the placebo group as compared to the control group retrieved from analyses of covariance (ANCOVA) on vection-induced fold changes of proteins (PFC) on day 2 (between-subject factors 'group' and 'sex', covariate PFC on day 1).

| Genes | Protein Accessions | *P*-value (Group) | No. of Peptides | Mean PFC day 1 (All) | Mean PFC day2 (Control) | Mean PFC day2 (Placebo) |
|---|---|---|---|---|---|---|
| **A2M** | P01023 | 0.000 | 633 | 0.100 | 0.052 | -0.033 |
| **ABCC9** | O60706 | 0.002 | 1 | 0.150 | -1.452 | 0.286 |
| **AHSG** | P02765 | 0.013 | 70 | 0.072 | -0.145 | 0.069 |
| **ALB;C3;APOA2** | P02768;P01024; P01781;P02652 | 0.035 | 1 | 0.579 | 0.833 | -0.433 |
| **ANKRD23** | Q86SG2 | 0.041 | 1 | 0.928 | -0.863 | 0.169 |
| **APOA1** | P02647 | 0.002 | 193 | 0.000 | -0.084 | 0.053 |
| **APOB** | P04114 | 0.000 | 1102 | 0.005 | 0.057 | -0.037 |
| **C19orf68** | Q86XI8 | 0.015 | 2 | 0.685 | -0.364 | 0.096 |
| **C1QC** | P02747 | 0.031 | 20 | 0.240 | 0.142 | -0.060 |
| **C3** | P01024 | 0.000 | 828 | 0.103 | 0.064 | -0.038 |
| **C4A** | P0C0L4 | 0.000 | 411 | 0.922 | 0.113 | -0.017 |
| **C4BPB** | P20851 | 0.044 | 6 | 0.837 | 0.324 | -0.272 |
| **C5** | P01031 | 0.020 | 158 | 0.178 | 0.106 | -0.020 |
| **CAT** | P04040 | 0.040 | 7 | 0.514 | 0.719 | -0.057 |
| **CCSER1** | Q9C0I3 | 0.028 | 1 | 0.975 | -0.252 | 0.222 |
| **CD163** | Q86VB7 | 0.006 | 2 | 0.077 | 1.145 | 0.019 |
| **CFH** | P08603 | 0.040 | 135 | 0.014 | 0.104 | -0.012 |
| **COG5** | Q9UP83 | 0.048 | 3 | 0.284 | -0.537 | 0.270 |
| **CP** | P00450 | 0.001 | 342 | 0.004 | 0.092 | -0.040 |
| **CRB1** | P82279 | 0.041 | 3 | 0.766 | -0.411 | 0.108 |
| **EVC** | P57679 | 0.020 | 1 | 0.003 | -0.738 | 0.698 |
| **F10** | P00742 | 0.003 | 15 | 0.954 | -0.415 | 0.190 |
| **FGA** | P02671 | 0.001 | 391 | 0.000 | 0.128 | -0.011 |
| **FGB** | P02675 | 0.000 | 313 | 0.030 | 0.048 | -0.086 |
| **FGG** | P02679 | 0.000 | 250 | 0.025 | 0.089 | -0.080 |
| **GAPDHS** | O14556 | 0.006 | 1 | 0.657 | -2.004 | 2.724 |
| **GC** | P02774 | 0.008 | 111 | 0.569 | 0.210 | 0.020 |
| **GYLTL1B** | Q8N3Y3 | 0.041 | 1 | 0.822 | -0.781 | 0.218 |
| **HBA1** | P69905 | 0.000 | 64 | 0.316 | 0.042 | -0.397 |
| **HBB** | P68871 | 0.001 | 47 | 0.005 | -0.071 | -0.357 |
| **HNRNPA1L2** | Q32P51 | 0.022 | 2 | 0.637 | 1.500 | -0.342 |
| **HP** | P00738 | 0.036 | 122 | 0.415 | 0.034 | -0.089 |
| **HPR;HP;GPLD1;FGB;GC;MAPT** | P00739;P00738; P80108;P02675; P02774;P10636 | 0.015 | 1 | 0.689 | 0.827 | -0.214 |
| **HPS3** | Q969F9 | 0.009 | 1 | 0.001 | -1.027 | 0.914 |
| **HSPA8** | P11142 | 0.030 | 10 | 0.821 | 0.472 | -0.311 |
| **IGFALS** | P35858 | 0.046 | 33 | 0.828 | 0.192 | -0.053 |
| **IGHG2** | P01859 | 0.032 | 85 | 0.043 | 0.120 | -0.032 |
| **IGHG3** | P01860 | 0.000 | 48 | 0.115 | 0.321 | -0.152 |

| | | | | | | |
|---|---|---|---|---|---|---|
| **IGHV3-23** | P01764 | 0.010 | 24 | 0.640 | 0.204 | -0.123 |
| **IGLC7** | A0M8Q6 | 0.029 | 4 | 0.404 | -0.558 | 0.306 |
| **JUP** | P14923 | 0.042 | 17 | 0.589 | 0.552 | -0.104 |
| **KIAA1614** | Q5VZ46 | 0.001 | 1 | 0.063 | -1.154 | 0.684 |
| **KRT10** | P13645 | 0.001 | 121 | 0.000 | -0.058 | 0.143 |
| **KRT6B;FGB** | P04259;P02675 | 0.013 | 1 | 0.955 | 0.576 | -0.246 |
| **LTBP1** | Q14766 | 0.042 | 4 | 0.102 | -0.189 | 0.226 |
| **LYZ** | P61626 | 0.044 | 7 | 0.696 | -0.381 | 0.377 |
| **MAPK8IP2** | Q13387 | 0.037 | 2 | 0.737 | -0.356 | 0.110 |
| **MYO18B** | Q8IUG5 | 0.005 | 1 | 0.645 | -0.417 | 0.959 |
| **MYO5B** | Q9ULV0 | 0.042 | 2 | 0.998 | 0.056 | -0.629 |
| **NKIRAS1** | Q9NYS0 | 0.016 | 1 | 0.137 | -1.563 | 1.074 |
| **ORM1** | P02763 | 0.016 | 38 | 0.878 | -0.182 | 0.110 |
| **PAPLN** | O95428 | 0.003 | 1 | 0.375 | 0.324 | -0.648 |
| **PDLIM1** | O00151 | 0.021 | 1 | 0.889 | 0.890 | -1.432 |
| **PFN2** | P35080 | 0.040 | 1 | 0.300 | 1.283 | -0.719 |
| **PIH1D2** | Q8WWB5 | 0.028 | 1 | 0.298 | 1.795 | 0.038 |
| **PPIA** | P62937 | 0.048 | 4 | 0.953 | 0.635 | -0.061 |
| **PRDX2** | P32119 | 0.033 | 11 | 0.037 | -0.025 | -0.515 |
| **PROC** | P04070 | 0.033 | 9 | 0.821 | -0.406 | 0.359 |
| **PRSS27** | Q9BQR3 | 0.028 | 2 | 0.062 | -0.907 | -0.117 |
| **PZP** | P20742 | 0.042 | 24 | 0.013 | 0.295 | -0.084 |
| **RAB3D** | O95716 | 0.021 | 1 | 0.243 | -0.899 | 0.121 |
| **RPL14** | P50914 | 0.022 | 1 | 0.162 | -2.392 | 0.323 |
| **SERPINB12** | Q96P63 | 0.024 | 5 | 0.118 | -0.296 | 0.370 |
| **SERPIND1** | P05546 | 0.000 | 83 | 0.717 | 0.159 | -0.118 |
| **SPG11** | Q96JI7 | 0.027 | 1 | 0.501 | 0.551 | -0.558 |
| **TF** | P02787 | 0.012 | 355 | 0.000 | 0.079 | -0.021 |
| **TSPYL1** | Q9H0U9 | 0.039 | 2 | 0.854 | -1.026 | 0.285 |
| **TTC36** | A6NLP5 | 0.004 | 2 | 0.384 | 1.557 | -0.521 |
| **VIL1** | P09327 | 0.046 | 1 | 0.389 | -0.107 | 0.870 |
| **VWA3A** | A6NCI4 | 0.045 | 1 | 0.498 | -0.453 | 0.476 |
| ***N/A*** | P01703 | 0.006 | 4 | 0.040 | -0.582 | 0.195 |
| ***N/A*** | P01607 | 0.033 | 3 | 0.725 | -0.369 | 0.416 |
| ***N/A*** | P01614 | 0.034 | 2 | 0.406 | -0.459 | 0.155 |
| ***N/A*** | P06888 | 0.040 | 5 | 0.805 | -0.509 | 0.224 |

**Supplemental Table 4:** Significant Gene enrichment (GO) terms and related proteins resulting from GO enrichment analyses on 74 proteins differentially regulated in the placebo group (for list of 74 proteins, see supplemental table 2).

| Gene enrichment (GO) terms | *P*-value (FDR-corr.) | Genes |
|---|---|---|
| platelet degranulation | 0.022 | A2M, APOA1, FGA, FGB, FGG, ORM1, AHSG, TF |
| positive regulation of substrate adhesion-dependent cell spreading | 0.008 | APOA1, FGA, FGB, FGG |
| vitamin transport | 0.047 | APOA1, GC |
| positive regulation of lipid storage | 0.022 | C3, APOB |
| hydrogen peroxide catabolic process | 0.010 | CAT, PRDX2, HBB, HBA1 |
| cellular protein metabolic process | 0.000 | CP, C3, APOA1, FGA, FGG, AHSG, TF, PROC, APOB, SERPIND1, C4A, IGFALS, LYZ, LTBP1 |
| post-translational protein modification | 0.000 | CP, C3, APOA1, FGA, FGG, AHSG, TF, PROC, APOB, SERPIND1, C4A, LTBP1 |
| blood coagulation, common pathway | 0.022 | F10, FGA |
| blood coagulation | 0.002 | F10, FGA, FGB, FGG, PROC, SERPIND1, C4BPB, HBB |
| induction of bacterial agglutination | 0.022 | FGA, FGB |
| positive regulation of peptide hormone secretion | 0.002 | FGA, FGB, FGG |
| cellular protein complex assembly | 0.002 | FGA, FGB, FGG |
| negative regulation of extrinsic apoptotic signaling pathway via death domain receptors | 0.005 | FGA, FGB, FGG |
| positive regulation of heterotypic cell-cell adhesion | 0.005 | FGA, FGB, FGG |
| positive regulation of vasoconstriction | 0.005 | FGA, FGB, FGG |
| blood coagulation, fibrin clot formation | 0.010 | FGA, FGB, FGG |
| positive regulation of exocytosis | 0.010 | FGA, FGB, FGG |
| protein polymerization | 0.010 | FGA, FGB, FGG |
| plasminogen activation | 0.022 | FGA, FGB, FGG |
| negative regulation of endothelial cell apoptotic process | 0.039 | FGA, FGB, FGG |
| toll-like receptor signaling pathway | 0.010 | FGA, FGB, FGG, APOB |
| platelet aggregation | 0.038 | FGA, FGB, FGG, HBB |
| positive regulation of protein secretion | 0.010 | FGA, FGB, FGG, PPIA |
| response to hydrogen peroxide | 0.004 | HP, CAT, HBB, HBA1 |
| cellular oxidant detoxification | 0.022 | HP, CAT, HBB, HBA1 |
| positive regulation of cell death | 0.018 | HP, HBB, HBA1 |
| acute-phase response | 0.038 | HP, ORM1, AHSG, CD163 |
| regulation of complement activation | 0.011 | IGLC7, C3, C5, IGHV3-23, IGHG2, IGHG3, C1QC, CFH, C4A, C4BPB |
| complement activation | 0.022 | IGLC7, C3, C5, IGHV3-23, IGHG2, IGHG3, C1QC, CFH, C4A |
| innate immune response | 0.047 | IGLC7, IGHV3-23, IGHG2, IGHG3, FGA, FGB, C1QC, C4A, C4BPB |
| protein heterooligomerization | 0.039 | JUP, HBB, HBA1 |
| negative regulation of endopeptidase activity | 0.001 | PAPLN, A2M, C3, C5, AHSG, SERPIND1, C4A, PZP, SERPINB12 |
| neutrophil degranulation | 0.005 | RAB3D, HP, C3, ORM1, AHSG, CAT, HSPA8, JUP, LYZ, PPIA, HBB, SERPINB12 |

**Supplemental Table 6**: Significant prediction of DAS-Nausea, DAS-MS, and DAS-NTT in the placebo and control groups by PFC of significantly enriched GO terms (see Supplemental Table 4).

| Gene enrichment (GO) term | DAS-Nau Ctrl | DAS-MS Ctrl | DAS-NTT Ctrl | DAS-Nau Plc | DAS-MS Plc | DAS-NTT Plc | Proteins |
|---|---|---|---|---|---|---|---|
| | ***P*-value (FDR-corrected)** | | | | | | |
| **Placebo group** | | | | | | | |
| acute-phase response | 0.140 | 0.781 | 0.336 | **0.003** | 0.771 | 0.297 | HP, ORM1, AHSG, CD163 |
| blood coagulation, common pathway | 0.103 | 0.560 | 0.400 | 0.807 | 0.331 | **0.023** | FGA |
| induction of bacterial agglutination | 0.178 | 0.515 | 0.268 | 0.943 | 0.321 | **0.015** | FGA, FGB |
| positive regulation of peptide hormone secretion | 0.241 | 0.665 | 0.370 | 0.896 | 0.312 | **0.038** | FGA, FGB, FGG |
| platelet aggregation | 0.308 | 0.821 | 0.383 | 0.710 | 0.238 | **0.020** | FGA, FGB, FGG, HBB |
| blood coagulation | 0.445 | 0.226 | 0.533 | 0.598 | 0.360 | **0.050** | FGA, FGB, FGG, SERPIND1, HBB |
| regulation of complement activation | 0.063 | 0.164 | 0.804 | 0.706 | 0.698 | **0.026** | C3, C5, IGHV3_23, IGHG2, IGHG3, C1QC, CFH, C4A |
| **Control group** | | | | | | | |
| positive regulation of substrate adhesion-dependent cell spreading | 0.413 | **0.000** | 0.492 | 0.952 | 0.706 | **0.019** | APOA1, FGA, FGB, FGG |
| protein heterooligomerization | **0.049** | **0.006** | 0.540 | 0.404 | 0.095 | 0.072 | HBB, HBA1 |
| positive regulation of cell death | 0.110 | **0.021** | 0.714 | 0.632 | 0.182 | 0.130 | HP, HBB, HBA1 |
| hydrogen peroxide catabolic process | 0.087 | **0.022** | 0.204 | 0.492 | 0.184 | 0.141 | CAT, HBB, HBA1 |

Abbreviations: DAS, day-adjusted scores; PFC, protein fold changes; Nau, nausea; Ctrl, control group; MS, motion sickness score; NTT, normo-to-tachy-ratio; Plc, placebo group.

**Supplemental Table 7:** Proteins for which a significant amount of variance could be explained by 'group', 'sex', 'DAS-MS', or by any of the interaction terms.

| Genes | Protein Accessions | Intercept | MS | Group | sex | MS X Group | MS X sex | Group X sex | MS X Group X sex |
|---|---|---|---|---|---|---|---|---|---|
| | | | | | ***P*-value** | | | | |
| **A1BG** | P04217 | 0.035 | **0.000** | 0.147 | 0.097 | **0.000** | **0.001** | 0.156 | **0.001** |
| **ABCA13** | Q86UQ4 | 0.562 | **0.000** | 0.490 | 0.624 | **0.000** | **0.000** | 0.674 | **0.001** |
| **ADAD1** | Q96M93 | 0.159 | **0.001** | 0.629 | 0.057 | **0.014** | **0.000** | 0.275 | **0.002** |
| **ALB;** | P02768;P01619 | 0.208 | **0.011** | 0.308 | 0.887 | **0.005** | **0.022** | 0.767 | **0.006** |
| **ALB;C3;APOB** | P01765;P02768;P01024;P04114 | 0.037 | **0.000** | 0.227 | 0.195 | **0.000** | **0.001** | 0.746 | **0.000** |
| **ALB;FGA;LYZ;VWF;LOXHD1;** | P02768;P02671;P61626;P04275;Q8IVV2;P04208 | 0.052 | **0.000** | 0.051 | 0.088 | **0.001** | **0.003** | 0.069 | **0.012** |
| **ALOX12B** | O75342 | 0.837 | 0.442 | 0.969 | 0.859 | 0.211 | 0.207 | 0.870 | **0.013** |
| **ANXA2** | P07355 | 0.399 | 0.778 | 0.848 | 0.156 | 0.388 | 0.160 | 0.665 | **0.017** |
| **ARHGAP10** | A1A4S6 | 0.035 | **0.000** | **0.016** | **0.020** | **0.001** | **0.005** | **0.048** | **0.017** |
| **ATR** | Q13535 | 0.836 | **0.015** | 0.663 | 0.931 | **0.013** | **0.030** | 0.783 | **0.024** |
| **AZGP1** | P25311 | 0.182 | **0.033** | 0.185 | 0.188 | 0.053 | **0.040** | 0.349 | **0.025** |
| **BIN2** | Q9UBW5 | 0.062 | 0.112 | 0.152 | **0.014** | **0.049** | 0.122 | 0.106 | **0.035** |
| **BLMH** | Q13867 | 0.103 | 0.112 | 0.162 | 0.073 | 0.313 | **0.044** | 0.210 | **0.037** |
| **BRMS1** | Q9HCU9 | 0.020 | **0.006** | 0.147 | 0.067 | **0.013** | **0.023** | 0.203 | **0.037** |
| **CAB39L** | Q9H9S4 | 0.713 | **0.001** | 0.903 | 0.440 | **0.004** | **0.013** | 0.414 | **0.043** |
| **CCDC160** | A6NGH7 | 0.052 | **0.013** | 0.070 | 0.148 | **0.021** | 0.062 | 0.180 | **0.044** |
| **CDC6;IGHM;DSP;EZR;** | P04220;Q99741;P01871;P15924;P15311;P01768 | 0.003 | **0.021** | **0.003** | **0.003** | 0.076 | **0.000** | **0.006** | **0.000** |
| **CEP290** | O15078 | 0.034 | **0.010** | 0.058 | **0.049** | **0.026** | **0.029** | 0.066 | **0.045** |
| **CFD** | P00746 | 0.239 | 0.476 | 0.068 | 0.303 | 0.567 | **0.016** | 0.053 | **0.045** |
| **CLEC11A** | Q9Y240 | 0.056 | 0.083 | **0.048** | **0.007** | **0.027** | 0.062 | **0.001** | **0.048** |
| **COLEC10** | Q9Y6Z7 | 0.298 | **0.020** | 0.544 | 0.535 | **0.033** | 0.054 | 0.448 | 0.051 |
| **CREBRF** | Q8IUR6 | 0.176 | 0.625 | 0.182 | **0.042** | 0.445 | 0.225 | 0.069 | 0.056 |
| **DDB1** | Q16531 | 0.750 | **0.050** | 0.932 | 0.860 | 0.101 | **0.037** | 0.776 | 0.059 |
| **DDX54** | Q8TDD1 | 0.063 | **0.006** | 0.159 | 0.187 | **0.003** | 0.050 | 0.161 | 0.060 |
| **DNHD1** | Q96M86 | 0.828 | 0.066 | 0.737 | 0.584 | 0.104 | **0.027** | 0.402 | 0.065 |
| **ENO1** | P06733 | 0.950 | 0.080 | 0.168 | 0.963 | **0.026** | 0.127 | 0.122 | 0.066 |
| **EPM2AIP1** | Q7L775 | 0.100 | 0.161 | **0.021** | 0.179 | 0.164 | 0.160 | 0.072 | 0.066 |
| **FBLN5** | Q9UBX5 | 0.828 | 0.927 | 0.698 | 0.260 | 0.834 | **0.020** | 0.320 | 0.073 |
| **FSTL1** | Q12841 | 0.001 | **0.013** | **0.014** | **0.018** | **0.026** | **0.046** | 0.059 | 0.075 |
| **FZD3** | Q9NPG1 | 0.019 | 0.114 | 0.105 | 0.208 | **0.014** | 0.226 | 0.657 | 0.083 |
| **GRXCR2** | A6NFK2 | 0.577 | 0.129 | 0.664 | 0.842 | **0.044** | 0.203 | 0.451 | 0.084 |
| **GSN** | P06396 | 0.404 | 0.060 | 0.421 | 0.137 | **0.047** | 0.111 | 0.055 | 0.084 |
| **GUCY1B2** | O75343 | 0.014 | **0.025** | 0.076 | 0.085 | **0.030** | 0.096 | 0.337 | 0.095 |
| **GYLTL1B** | Q8N3Y3 | 0.211 | 0.095 | 0.283 | 0.668 | **0.019** | 0.226 | 0.483 | 0.100 |
| **HPS3** | Q969F9 | 0.479 | **0.008** | 0.722 | 0.776 | **0.035** | **0.043** | 0.794 | 0.109 |
| **HRG** | P04196 | 0.060 | **0.024** | 0.118 | 0.200 | **0.019** | 0.096 | 0.331 | 0.119 |
| **HSD3B1** | P14060 | 0.067 | **0.003** | 0.066 | 0.867 | **0.007** | 0.124 | 0.755 | 0.125 |
| **HTT** | P42858 | 0.379 | **0.023** | 0.291 | 0.452 | 0.083 | **0.010** | 0.260 | 0.127 |
| **IGFALS** | P35858 | 0.949 | 0.111 | 0.797 | 0.854 | 0.274 | **0.040** | 0.958 | 0.128 |

| Genes | Protein Accessions | Intercept | MS | Group | sex | MS X Group | MS X sex | Group X sex | MS X Group X sex |
|---|---|---|---|---|---|---|---|---|---|
| **IGHG2** | P01859 | 0.766 | 0.105 | 0.416 | 0.230 | **0.037** | 0.680 | 0.132 | 0.130 |
| **IGHM;** | P01871;P01773 | 0.909 | 0.703 | 0.531 | 0.073 | 0.667 | 0.083 | **0.022** | 0.139 |
| **IZUMO4** | Q1ZYL8 | 0.088 | **0.031** | 0.073 | 0.061 | **0.014** | 0.123 | **0.022** | 0.142 |
| **KIF14** | Q15058 | 0.144 | **0.031** | 0.138 | 0.515 | **0.009** | 0.117 | 0.253 | 0.143 |
| **KIF17** | Q9P2E2 | 0.109 | 0.056 | 0.334 | 0.219 | **0.049** | 0.341 | 0.422 | 0.146 |
| **KRT10** | P13645 | 0.900 | **0.041** | 0.337 | 0.955 | 0.100 | **0.034** | 0.341 | 0.153 |
| **KRT13** | P13646 | 0.903 | 0.955 | 0.983 | **0.047** | 0.909 | 0.131 | 0.152 | 0.209 |
| **KRT6B** | P04259 | 0.281 | 0.587 | 0.243 | 0.063 | 0.590 | 0.274 | **0.015** | 0.225 |
| **KRT9** | P35527 | 0.853 | 0.453 | 0.297 | 0.429 | 0.886 | 0.082 | **0.047** | 0.226 |
| **MAPT** | P10636 | 0.215 | 0.094 | 0.785 | **0.004** | 0.567 | **0.035** | 0.103 | 0.234 |
| **MYO5B** | Q9ULV0 | 0.088 | 0.278 | 0.103 | **0.048** | 0.220 | 0.337 | 0.051 | 0.235 |
| **NDUFS2** | O75306 | 0.085 | 0.287 | **0.034** | 0.090 | 0.402 | 0.254 | **0.030** | 0.252 |
| **NRIP1** | P48552 | 0.647 | **0.050** | 0.723 | 0.723 | **0.025** | 0.251 | 0.358 | 0.253 |
| **NRXN1** | Q9ULB1 | 0.484 | 0.488 | 0.288 | 0.063 | 0.281 | 0.270 | **0.020** | 0.254 |
| **OSBPL1A** | Q9BXW6 | 0.895 | **0.036** | 0.987 | 0.674 | **0.030** | 0.198 | 0.928 | 0.257 |
| **OSBPL5** | Q9H0X9 | 0.695 | **0.038** | 0.272 | 0.470 | 0.232 | 0.171 | 0.955 | 0.266 |
| **PHACTR1** | Q9C0D0 | 0.309 | 0.220 | 0.242 | 0.119 | 0.571 | 0.118 | **0.020** | 0.275 |
| **PLCXD1** | Q9NUJ7 | 0.119 | **0.017** | 0.371 | 0.178 | 0.055 | **0.048** | 0.312 | 0.288 |
| **PLG** | P00747 | 0.377 | **0.041** | 0.621 | 0.512 | **0.037** | 0.475 | 0.571 | 0.301 |
| **POLI** | Q9UNA4 | 0.131 | 0.173 | **0.049** | 0.513 | 0.117 | 0.618 | 0.220 | 0.324 |
| **PPIP5K2** | O43314 | 0.116 | 0.222 | **0.015** | 0.473 | 0.288 | 0.266 | 0.149 | 0.334 |
| **PPP4R3CP** | Q6ZMV5 | 0.349 | **0.014** | 0.238 | 0.693 | 0.081 | 0.059 | 0.296 | 0.334 |
| **RICTOR** | Q6R327 | 0.065 | 0.967 | **0.045** | **0.040** | 0.835 | 0.871 | **0.012** | 0.336 |
| **RNH1** | P13489 | 0.062 | 0.498 | 0.084 | 0.111 | 0.273 | 0.682 | **0.042** | 0.391 |
| **S100A7** | P31151 | 0.427 | 0.263 | 0.180 | 0.263 | 0.300 | 0.324 | **0.047** | 0.397 |
| **S100A9** | P06702 | 0.019 | 0.096 | **0.048** | **0.037** | 0.139 | 0.372 | 0.069 | 0.454 |
| **SERPINB12** | Q96P63 | 0.864 | **0.038** | 0.777 | 0.568 | 0.089 | 0.245 | 0.515 | 0.462 |
| **SERPINF1** | P36955 | 0.149 | 0.671 | 0.112 | 0.097 | 0.576 | 0.704 | **0.032** | 0.503 |
| **SGSM1** | Q2NKQ1 | 0.102 | 0.167 | 0.066 | 0.100 | 0.162 | 0.375 | **0.039** | 0.513 |
| **SHANK1** | Q9Y566 | 0.240 | **0.018** | 0.191 | 0.873 | **0.022** | 0.241 | 0.409 | 0.513 |
| **SKAP1** | Q86WV1 | 0.691 | 0.697 | 0.779 | 0.054 | 0.723 | 0.175 | **0.020** | 0.548 |
| **SLC35C2** | Q9NQQ7 | 0.129 | **0.036** | 0.153 | 0.742 | 0.200 | 0.206 | 0.861 | 0.600 |
| **SORCS2** | Q96PQ0 | 0.599 | 0.435 | 0.649 | 0.098 | 0.638 | 0.376 | **0.037** | 0.645 |
| **SPEN** | Q96T58 | 0.140 | 0.817 | 0.343 | **0.042** | 0.840 | 0.624 | 0.204 | 0.681 |
| **SPP1** | P10451 | 0.358 | 0.054 | 0.376 | 0.618 | **0.033** | 0.343 | 0.363 | 0.724 |
| **SVOPL** | Q8N434 | 0.137 | **0.030** | 0.094 | 0.788 | 0.167 | 0.293 | 0.778 | 0.749 |
| **SYNE1** | Q8NF91 | 0.874 | **0.009** | 0.500 | 0.164 | 0.056 | 0.217 | 0.075 | 0.764 |
| **TRAP1** | Q12931 | 0.970 | 0.901 | 0.777 | 0.342 | 0.955 | **0.046** | 0.626 | 0.772 |
| **TXN** | P10599 | 0.193 | 0.915 | 0.388 | **0.044** | 0.857 | 0.552 | 0.071 | 0.794 |
| **VCAM1** | P19320 | 0.307 | **0.018** | 0.264 | 0.950 | 0.331 | 0.084 | 0.959 | 0.835 |
| **ZG16B** | Q96DA0 | 0.447 | 0.752 | 0.229 | 0.174 | 0.996 | 0.874 | **0.025** | 0.856 |
| **ZNF438** | Q7Z4V0 | 0.564 | **0.021** | 0.316 | 0.484 | 0.086 | 0.465 | 0.906 | 0.866 |
|  | P01596 | 0.058 | 0.901 | **0.007** | 0.256 | 0.982 | 0.982 | **0.036** | 0.868 |
|  | P01762 | 0.056 | 0.087 | **0.027** | 0.353 | 0.437 | 0.321 | 0.317 | 0.924 |

| Genes | Protein Accessions | Intercept | MS | Group | sex | MS X Group | MS X sex | Group X sex | MS X Group X sex |
|---|---|---|---|---|---|---|---|---|---|
| | P80422 | 0.028 | 0.119 | **0.008** | 0.384 | 0.300 | 0.508 | 0.171 | 0.982 |
| | P01707 | 0.211 | 0.789 | 0.127 | 0.151 | 0.952 | 0.388 | **0.028** | 0.996 |

Abbreviations: DAS-MS, day-adjusted scores of motion sickness.

**Supplemental Table 8:** Proteins for which a significant amount of variance could be explained by ‘group’, ‘sex’, ‘DAS-NTT’, or by any of the interaction terms.

| Genes | PG Protein Accessions | Intercept | NTT | Group | sex | NTT X Group | NTT X Sex | Group X Sex | NTT X Group X sex |
|---|---|---|---|---|---|---|---|---|---|
| | | *P*-Value | | | | | | | |
| **ABCA13** | Q86UQ4 | 0.810 | 0.497 | 0.677 | 0.053 | 0.694 | 0.051 | **0.013** | 0.704 |
| **ABCA6** | Q8N139 | 0.439 | 0.242 | **0.015** | 0.533 | 0.994 | 0.270 | **0.048** | 0.984 |
| **ABCC9** | O60706 | 0.045 | 0.191 | **0.037** | 0.329 | 0.181 | **0.044** | 0.807 | **0.013** |
| **ACTN2** | P35609 | 0.018 | 0.220 | 0.076 | **0.012** | 0.197 | 0.444 | **0.029** | 0.132 |
| **ALB;VNN1; FGG; TECTA** | P02768;O95497;P02679; P01597;O75443 | 0.255 | 0.951 | **0.019** | 0.748 | 0.311 | 0.876 | 0.180 | 0.208 |
| **ANKRD11** | Q6UB99 | 0.898 | **0.017** | 0.934 | 0.601 | **0.040** | 0.124 | 0.553 | 0.391 |
| **ANKRD55** | Q3KP44 | 0.002 | 0.225 | **0.002** | **0.002** | 0.436 | 0.413 | **0.007** | 0.456 |
| **APOC4** | P55056 | 0.011 | **0.024** | **0.032** | **0.004** | 0.189 | 0.196 | **0.015** | 0.555 |
| **APOM** | O95445 | 0.213 | 0.713 | 0.378 | 0.238 | 0.769 | 0.573 | **0.044** | 0.347 |
| **APP** | P05067 | 0.025 | 0.927 | **0.008** | **0.009** | 0.444 | 0.905 | **0.007** | 0.677 |
| **ATAD2** | Q6PL18 | 0.276 | **0.041** | 0.263 | 0.758 | 0.157 | 0.123 | 0.612 | 0.116 |
| **AZGP1** | P25311 | 0.038 | 0.621 | 0.064 | 0.071 | 0.640 | 0.621 | **0.023** | 0.540 |
| **B3GNT4** | Q9C0J1 | 0.490 | **0.045** | 0.339 | 0.550 | **0.037** | **0.016** | 0.223 | **0.040** |
| **BIN2** | Q9UBW5 | 0.033 | 0.100 | **0.045** | 0.232 | 0.639 | 0.258 | 0.475 | 0.998 |
| **BLMH** | Q13867 | 0.128 | 0.808 | 0.237 | **0.042** | 0.951 | 0.783 | 0.162 | 0.463 |
| **C5;APOA4** | P01031;P01766; P06727 | 0.050 | **0.009** | 0.265 | **0.021** | **0.002** | 0.057 | 0.069 | 0.051 |
| **C5orf42** | Q9H799 | 0.092 | 0.246 | 0.085 | 0.064 | 0.326 | 0.496 | **0.044** | 0.819 |
| **CAB39L** | Q9H9S4 | 0.084 | 0.610 | 0.102 | **0.046** | 0.715 | 0.658 | **0.044** | 0.659 |
| **CD44** | P16070 | 0.732 | 0.511 | 0.963 | 0.989 | 0.549 | **0.033** | 0.818 | 0.151 |
| **CFHR2** | P36980 | 0.071 | 0.406 | **0.026** | 0.113 | 0.950 | 0.405 | **0.042** | 0.909 |
| **CNTNAP4** | Q9C0A0 | 0.331 | 0.071 | 0.295 | 0.728 | 0.201 | **0.006** | 0.659 | **0.022** |
| **COL1A2** | P08123;P01619 | 0.992 | 0.239 | 0.967 | 0.604 | 0.073 | 0.196 | 0.520 | **0.045** |
| **COL6A3** | P12111 | 0.954 | 0.080 | 0.639 | 0.431 | 0.052 | 0.138 | 0.150 | **0.024** |
| **CP** | P00450 | 0.106 | 0.696 | 0.096 | 0.275 | 0.927 | 0.744 | **0.040** | 0.795 |
| **CPA4** | Q9UI42 | 0.696 | 0.838 | 0.986 | **0.042** | 0.473 | 0.465 | 0.095 | 0.219 |
| **CPN2** | P22792 | 0.310 | 0.088 | 0.899 | 0.381 | **0.025** | 0.097 | 0.907 | 0.082 |
| **DAAM1** | Q9Y4D1 | 0.106 | 0.576 | 0.093 | 0.223 | 0.567 | 0.500 | **0.032** | 0.164 |
| **DCD** | P81605 | 0.113 | 0.186 | 0.087 | 0.124 | 0.252 | 0.227 | **0.039** | 0.259 |
| **DDX54** | Q8TDD1 | 0.604 | 0.932 | 0.404 | **0.046** | 0.452 | 0.233 | 0.702 | 0.246 |
| **DNHD1** | Q96M86 | 0.985 | **0.001** | 0.620 | 0.855 | **0.005** | **0.006** | 0.723 | **0.028** |
| **DROSHA** | Q9NRR4 | 0.014 | 0.062 | 0.054 | **0.021** | 0.085 | 0.191 | **0.048** | 0.419 |
| **EDEM3** | Q9BZQ6 | 0.977 | **0.000** | 0.513 | 0.926 | **0.000** | **0.003** | 0.734 | **0.002** |
| **ENO1** | P06733 | 0.019 | 0.438 | **0.013** | 0.246 | 0.420 | 0.460 | 0.212 | 0.423 |
| **F2** | P00734 | 0.424 | 0.923 | 0.116 | 0.165 | 0.878 | 0.766 | **0.023** | 0.488 |
| **FGA** | P02671 | 0.137 | 0.470 | 0.133 | 0.179 | 0.631 | 0.335 | **0.034** | 0.573 |
| **FGB** | P02675;P01597 | 0.898 | 0.321 | 0.909 | 0.135 | 0.257 | **0.049** | 0.143 | 0.137 |
| **GAB2** | Q9UQC2 | 0.536 | 0.061 | 0.961 | 0.376 | **0.023** | **0.046** | 0.834 | 0.140 |
| **GAPDHS** | O14556 | 0.843 | **0.017** | 0.945 | 0.274 | 0.051 | 0.425 | 0.516 | 0.236 |
| **GRXCR2** | A6NFK2 | 0.016 | 0.108 | 0.057 | **0.046** | 0.124 | 0.112 | 0.204 | 0.095 |
| **GUCY1B2** | O75343 | 0.604 | 0.116 | 0.258 | 0.664 | 0.379 | **0.024** | 0.112 | 0.215 |

| Genes | PG Protein Accessions | Intercept | NTT | Group | sex | NTT X Group | NTT X Sex | Group X Sex | NTT X Group X sex |
|---|---|---|---|---|---|---|---|---|---|
| **HEPHL1** | Q6MZM0 | 0.923 | **0.015** | 0.610 | 0.371 | 0.633 | **0.015** | 0.472 | 0.611 |
| **HUWE1** | Q7Z6Z7 | 0.413 | 0.520 | 0.823 | 0.370 | 0.443 | 0.151 | 0.571 | **0.026** |
| **IGFALS** | P35858 | 0.068 | 0.739 | 0.099 | **0.048** | 0.632 | 0.532 | **0.026** | 0.451 |
| **IGHM** | P01871;P01773 | 0.132 | 0.255 | **0.007** | 0.270 | 0.829 | 0.269 | **0.035** | 0.495 |
| **IGHV1OR21-1** | A6NJS3 | 0.029 | 0.090 | 0.078 | 0.053 | **0.036** | 0.159 | 0.169 | 0.166 |
| **KIF14** | Q15058 | 0.613 | 0.973 | 0.564 | 0.920 | 0.246 | 0.138 | 0.948 | **0.024** |
| **KRT23** | Q9C075 | 0.088 | 0.457 | 0.076 | **0.021** | 0.546 | 0.268 | **0.023** | 0.444 |
| **KRT6B** | P04259 | 0.117 | 0.441 | **0.022** | 0.921 | 0.319 | 0.071 | 0.444 | 0.230 |
| **KRT77** | Q7Z794 | 0.247 | 0.795 | 0.106 | 0.172 | 0.966 | 0.375 | **0.035** | 0.537 |
| **KRT9** | P35527 | 0.416 | 0.782 | 0.135 | 0.259 | 0.869 | 0.549 | **0.029** | 0.687 |
| **LTN1** | O94822 | 0.101 | 0.619 | **0.024** | 0.125 | 0.641 | 0.414 | 0.096 | 0.661 |
| **LYVE1** | Q9Y5Y7 | 0.414 | 0.662 | 0.434 | 0.476 | 0.605 | **0.032** | 0.686 | **0.021** |
| **MED30** | Q96HR3 | 0.384 | **0.040** | 0.723 | 0.136 | **0.046** | **0.025** | 0.299 | **0.035** |
| **MLPH** | Q9BV36 | 0.089 | 0.053 | **0.030** | 0.175 | 0.094 | 0.144 | 0.064 | 0.249 |
| **MMP25** | Q9NPA2 | 0.658 | 0.183 | 0.395 | 0.583 | 0.077 | 0.086 | 0.465 | **0.047** |
| **MYO5B** | Q9ULV0 | 0.023 | 0.844 | **0.006** | 0.169 | 0.510 | 0.786 | 0.167 | 0.665 |
| **NKIRAS1** | Q9NYS0 | 0.013 | 0.371 | **0.004** | 0.055 | 0.847 | 0.861 | **0.050** | 0.373 |
| **NR0B1** | P51843 | 0.995 | 0.077 | 0.646 | 0.665 | **0.046** | 0.124 | 0.822 | 0.096 |
| **NRIP1** | P48552 | 0.344 | 0.877 | 0.075 | 0.831 | 0.613 | **0.039** | 0.171 | **0.040** |
| **NRXN1** | Q9ULB1 | 0.021 | 0.129 | **0.015** | **0.041** | 0.235 | 0.279 | 0.161 | 0.718 |
| **OSBPL1A** | Q9BXW6 | 0.200 | **0.004** | 0.273 | 0.170 | **0.014** | **0.012** | 0.224 | **0.035** |
| **PFN2** | P35080 | 0.084 | 0.699 | **0.038** | 0.400 | 0.611 | 0.256 | 0.138 | 0.112 |
| **PHACTR1** | Q9C0D0 | 0.148 | 0.800 | 0.075 | 0.058 | 0.948 | 0.886 | **0.022** | 0.840 |
| **PLG** | P00747 | 0.602 | 0.712 | 0.415 | 0.188 | 0.965 | 0.144 | **0.034** | 0.600 |
| **PRDX6** | P30041 | 0.612 | 0.072 | 0.836 | 0.955 | **0.017** | 0.179 | 0.628 | 0.153 |
| **QSOX1** | O00391 | 0.080 | 0.623 | 0.048 | 0.214 | 0.110 | 0.422 | 0.685 | **0.006** |
| **RBM44** | Q6ZP01 | 0.079 | 0.079 | 0.127 | 0.617 | **0.034** | 0.221 | 0.684 | 0.079 |
| **RELN** | P78509 | 0.430 | 0.987 | 0.552 | 0.691 | 0.727 | 0.200 | 0.488 | **0.047** |
| **RIC8B** | Q9NVN3 | 0.120 | **0.024** | **0.031** | 0.166 | **0.028** | 0.054 | **0.025** | **0.024** |
| **RPL14** | P50914 | 0.390 | 0.488 | 0.911 | **0.012** | 0.573 | 0.708 | 0.168 | 0.911 |
| **SERPINA1** | P01009 | 0.085 | 0.302 | 0.163 | 0.097 | 0.356 | 0.394 | **0.033** | 0.354 |
| **SERPINF1** | P36955 | 0.032 | 0.324 | **0.031** | **0.013** | 0.587 | 0.243 | **0.006** | 0.176 |
| **SKAP1** | Q86WV1 | 0.123 | 0.204 | **0.029** | **0.027** | 0.450 | 0.105 | **0.002** | 0.360 |
| **SLK** | Q9H2G2 | 0.561 | 0.428 | 0.741 | 0.820 | **0.037** | 0.324 | 0.740 | **0.009** |
| **SORCS2** | Q96PQ0 | 0.023 | 0.315 | **0.001** | 0.173 | 0.756 | 0.190 | **0.014** | 0.446 |
| **SOS2** | Q07890 | 0.645 | **0.018** | 0.755 | 0.913 | **0.020** | 0.113 | 0.315 | 0.177 |
| **SPARC** | P09486 | 0.074 | 0.869 | **0.044** | 0.183 | 0.510 | 0.996 | 0.151 | 0.667 |
| **SPTAN1** | Q13813 | 0.171 | 0.511 | 0.136 | **0.043** | 0.983 | 0.807 | **0.015** | 0.711 |
| **TEX11** | Q8IYF3 | 0.999 | 0.065 | 0.830 | 0.801 | 0.093 | 0.111 | 0.808 | **0.038** |
| **TGM1** | P22735 | 0.218 | **0.004** | 0.523 | **0.034** | **0.006** | 0.064 | 0.143 | 0.126 |
| **TGM3** | Q08188 | 0.053 | 0.187 | **0.025** | 0.111 | 0.370 | 0.279 | 0.080 | 0.656 |
| **TRAP1** | Q12931 | 0.039 | 0.056 | **0.013** | 0.187 | 0.307 | 0.454 | 0.097 | 0.969 |
| **TSC1** | Q92574 | 0.001 | 0.065 | **0.002** | **0.004** | 0.180 | 0.084 | **0.001** | 0.259 |
| **TTC36** | A6NLP5 | 0.039 | 0.393 | **0.022** | 0.179 | 0.412 | 0.670 | 0.176 | 0.867 |

| Genes | PG Protein Accessions | Intercept | NTT | Group | sex | NTT X Group | NTT X Sex | Group X Sex | NTT X Group X sex |
|---|---|---|---|---|---|---|---|---|---|
| **TXN** | P10599 | 0.111 | **0.001** | 0.126 | 0.243 | **0.006** | **0.003** | 0.595 | 0.133 |
| **ZG16B** | Q96DA0 | 0.255 | 0.673 | 0.074 | 0.193 | 0.834 | 0.605 | **0.035** | 0.663 |
| **ZNFX1** | Q9P2E3 | 0.448 | 0.989 | 0.254 | 0.817 | 0.484 | 0.109 | 0.223 | **0.028** |
| | P01597 | 0.048 | 0.079 | **0.048** | **0.048** | 0.347 | 0.271 | 0.088 | 0.859 |
| | P01772 | 0.771 | 0.136 | 0.730 | 0.751 | 0.076 | 0.122 | 0.899 | **0.033** |
| | P01778 | 0.389 | 0.235 | 0.377 | 0.577 | 0.158 | 0.094 | 0.473 | **0.049** |
| | P80422 | 0.395 | 0.786 | 0.327 | 0.111 | 0.506 | 0.756 | **0.031** | 0.864 |
| | P01619 | 0.942 | 0.944 | 0.942 | 0.230 | 0.946 | **0.042** | 0.312 | **0.042** |

Abbreviations: DAS-NTT, day-adjusted scores of normo-to-tachy ratio.

**Supplemental Table 9**: Enriched GO groups based on proteins for which a significant amount of variance could be explained by the factors 'group', 'sex', by 'DAS-Nausea', or by any of the interaction terms.

| *P*-value | GO group | Genes |
|---|---|---|
| 0.000 | cholesterol metabolic process | APOL1 APOA2 APOB APP PON1 OSBPL1A |
| 0.000 | cell adhesion | COL6A5 FN1 APP GP1BA AZGP1 ACTN2 IGFALS PKP1 POSTN GRHL2 |
| 0.001 | protein localization to plasma membrane | ANXA2 FLNA ACTN2 SKAP1 |
| 0.002 | mating behavior | APP MAPK8IP2 |
| 0.004 | response to insulin | EIF6 EPM2AIP1 TSC1 |
| 0.005 | regulation of translation | APP TSC1 |
| 0.005 | fertilization | APOB TEX11 |
| 0.005 | positive regulation of stress-activated MAPK cascade | MAPK8IP2 CARD9 |
| 0.005 | activation of GTPase activity | SGSM1 TSC1 |
| 0.008 | neural tube closure | GRHL2 TSC1 FZD3 |
| 0.009 | regulation of protein phosphorylation | FN1 HSPB1 |
| 0.009 | hair follicle development | FZD3 KRT84 |
| 0.009 | protein O-linked glycosylation | PLOD2 POMT2 |
| 0.009 | positive regulation of peptidase activity | FN1 APP |
| 0.009 | cell junction assembly | FLNA GRHL2 |
| 0.009 | sodium ion transmembrane transport | GRIK4 SCN2A |
| 0.009 | synapse organization | APP TSC1 |
| 0.009 | regulation of NMDA receptor activity | APP MAPK8IP2 |
| 0.013 | post-translational protein modification | APOL1 SERPINC1 APOA2 FN1 APOB APP |
| 0.015 | adult locomotory behavior | APP TSC1 |
| 0.015 | social behavior | MAPK8IP2 NRXN1 |
| 0.015 | ionotropic glutamate receptor signaling pathway | APP GRIK4 |
| 0.015 | positive regulation of JNK cascade | APP CARD9 |
| 0.015 | neuron apoptotic process | APP SCN2A |
| 0.015 | excitatory postsynaptic potential | MAPK8IP2 GRIK4 |
| 0.015 | cellular response to retinoic acid | KRT13 SERPINF1 |
| 0.017 | cytoskeleton organization | KRT6B KRT16 KRT13 KRT84 |
| 0.020 | cornification | KRT6B KRT16 KRT13 FLG PKP1 KRT84 |
| 0.021 | negative regulation of protein kinase activity | HSPB1 GP1BA |
| 0.021 | myelination | TSC1 SCN2A |
| 0.023 | cellular protein metabolic process | APOL1 SERPINC1 APOA2 FN1 APOB APP IGFALS |
| 0.029 | positive regulation of gene expression | FN1 APOB APP PKP1 |
| 0.029 | I-kappaB kinase/NF-kappaB signaling | CARD9 NKIRAS1 |
| 0.029 | learning | APP NRXN1 |
| 0.029 | chylomicron remodeling | APOA2 APOB |
| 0.029 | very-low-density lipoprotein particle assembly | ACSL3 APOB |
| 0.029 | negative regulation of catalytic activity | ANXA2 PHACTR1 |

| *P*-value | GO group | Genes |
|---:|---|---|
| 0.029 | establishment of skin barrier | KRT16 FLG |
| 0.030 | platelet aggregation | HSPB1 GP1BA FLNA |
| 0.033 | keratinization | KRT6B KRT16 KRT13 PKP1 KRT84 |
| 0.037 | kidney development | SERPINF1 TSC1 |
| 0.037 | low-density lipoprotein particle remodeling | APOA2 APOB |
| 0.037 | chylomicron assembly | APOA2 APOB |
| 0.037 | neuromuscular process controlling balance | APP NRXN1 |
| 0.045 | vesicle-mediated transport | OSBPL1A MYO5B |
| 0.045 | actin cytoskeleton reorganization | FLNA PHACTR1 |
| 0.045 | positive regulation of fibroblast proliferation | FN1 ANXA2 |
| 0.045 | protein heterooligomerization | FGFRL1 TSC1 |
| 0.048 | retina homeostasis | HSPB1 AZGP1 ZG16B |

Note: Significant GO groups with single genes were omitted.
Abbreviations: DAS-Nausea, day-adjusted scores of nausea.

**Supplemental Table 10:** Enriched groups of proteins for which a significant amount of variance could be explained by the factors 'group', 'sex', by 'DAS-MS', or by any of the interaction terms.

| *P*-value | GO group | Genes |
|---|---|---|
| 0.002 | adult behavior | NRXN1 SHANK1 |
| 0.002 | vocal learning | HTT NRXN1 |
| 0.002 | negative regulation of catalytic activity | ANXA2 RNH1 PHACTR1 |
| 0.004 | vesicle-mediated transport | OSBPL1A KIF17 MYO5B |
| 0.004 | neutrophil degranulation | CEP290 CFD A1BG GSN S100A9 ANXA2 S100A7 ABCA13 SERPINB12 BIN2 |
| 0.004 | protein destabilization | GSN HTT |
| 0.004 | vocalization behavior | NRXN1 SHANK1 |
| 0.004 | amyloid fibril formation | GSN MAPT |
| 0.009 | positive regulation of neurogenesis | SERPINF1 SPEN |
| 0.009 | interaction with symbiont | PLG DDB1 |
| 0.009 | positive regulation of excitatory postsynaptic potential | NRXN1 SHANK1 |
| 0.012 | transcription, DNA-templated | ENO1 TXN NRIP1 ZNF438 DDX54 SPEN BRMS1 |
| 0.014 | activation of cysteine-type endopeptidase activity involved in apoptotic process | S100A9 MAPT |
| 0.014 | social behavior | NRXN1 SHANK1 |
| 0.014 | tissue regeneration | PLG GSN |
| 0.014 | cellular response to retinoic acid | KRT13 SERPINF1 |
| 0.014 | negative regulation of gene expression | MAPT SERPINF1 SLC35C2 |
| 0.017 | fibrinolysis | PLG HRG ANXA2 |
| 0.017 | antimicrobial humoral immune response mediated by antimicrobial peptide | HRG S100A9 S100A7 |
| 0.018 | negative regulation of transcription from RNA polymerase II promoter | ENO1 TXN NRIP1 SPEN BRMS1 |
| 0.020 | response to radiation | TXN KRT13 |
| 0.027 | negative regulation of fibrinolysis | PLG HRG |
| 0.030 | positive regulation of transcription, DNA-templated | CEP290 SPP1 NRIP1 SKAP1 |
| 0.031 | DNA repair | ATR DDB1 POLI |
| 0.034 | RNA processing | DDX54 ADAD1 |
| 0.034 | peptidyl-serine phosphorylation | ATR RICTOR |
| 0.034 | negative regulation of cell growth | HRG ENO1 |
| 0.034 | neuromuscular process controlling balance | NRXN1 SHANK1 |
| 0.042 | regulation of gene expression | HRG RICTOR |
| 0.042 | actin cytoskeleton reorganization | RICTOR PHACTR1 |
| 0.044 | aging | GSN VCAM1 SERPINF1 |
| 0.048 | extracellular matrix disassembly | PLG GSN SPP1 |

Note: Significant GO groups with single genes were omitted.
Abbreviations: DAS-MS, day-adjusted scores of motion sickness.

**Supplemental Table 11:** Enriched groups of proteins for which a significant amount of variance could be explained by the factors 'group', 'sex', by 'DAS-NTT, or by any of the interaction terms.

| *P*-value | GO group | Genes |
|---|---|---|
| 0.001 | positive regulation of peptidyl-serine phosphorylation | APP TXN CD44 PFN2 |
| 0.002 | regulation of grooming behavior | CNTNAP4 NRXN1 |
| 0.002 | copper ion transport | CP HEPHL1 |
| 0.002 | positive regulation of small GTPase mediated signal transduction | RELN SOS2 |
| 0.002 | positive regulation of synaptic transmission, glutamatergic | RELN NRXN1 |
| 0.002 | positive regulation of synapse maturation | RELN NRXN1 |
| 0.002 | postsynaptic density protein 95 clustering | RELN NRXN1 |
| 0.002 | hippocampus development | RELN KIF14 TSC1 |
| 0.003 | axon guidance | RELN SPTAN1 NRXN1 GAB2 |
| 0.004 | kidney development | SERPINF1 TSC1 C5orf42 |
| 0.005 | cell redox homeostasis | QSOX1 TXN PRDX6 |
| 0.006 | regulation of translation | APP TSC1 |
| 0.006 | dendrite development | APP RELN |
| 0.006 | cell envelope organization | TGM1 TGM3 |
| 0.006 | positive regulation of long-term synaptic potentiation | APP RELN |
| 0.011 | potassium ion transport | ABCC9 TSC1 |
| 0.011 | response to lead ion | APP SPARC |
| 0.011 | negative regulation of macroautophagy | QSOX1 TSC1 |
| 0.011 | positive regulation of heterotypic cell-cell adhesion | FGA CD44 |
| 0.011 | synapse organization | APP TSC1 |
| 0.011 | bone development | SPARC ANKRD11 |
| 0.011 | regulation of NMDA receptor activity | APP RELN |
| 0.011 | positive regulation of excitatory postsynaptic potential | RELN NRXN1 |
| 0.012 | cellular protein metabolic process | QSOX1 CP F2 PLG SERPINA1 FGA APP IGFALS |
| 0.016 | platelet degranulation | QSOX1 PLG SERPINA1 FGA APP SPARC ACTN2 |
| 0.017 | adult locomotory behavior | APP TSC1 |
| 0.017 | hyaluronan catabolic process | CD44 LYVE1 |
| 0.017 | androgen receptor signaling pathway | NRIP1 MED30 |
| 0.018 | cell-matrix adhesion | FGA CD44 TSC1 LYVE1 |
| 0.019 | cellular oxidant detoxification | APOM TXN PRDX6 |
| 0.021 | neutrophil degranulation | QSOX1 SERPINA1 CD44 PRDX6 SPTAN1 HUWE1 ABCA13 MMP25 BIN2 |
| 0.022 | cell adhesion | APP COL6A3 AZGP1 ACTN2 IGFALS RELN CNTNAP4 |
| 0.022 | fibrinolysis | F2 PLG FGA |
| 0.023 | transmembrane transport | ABCC9 AZGP1 ABCA13 ABCA6 |

| | | |
|---|---|---|
| 0.024 | transcription initiation from RNA polymerase II promoter | NR0B1 MED30 |
| 0.024 | activation of protein kinase activity | KIF14 SLK |
| 0.024 | regulation of molecular function | ABCC9 NRXN1 |
| 0.031 | lipid transport | APOC4 ABCA13 ABCA6 |
| 0.031 | protein stabilization | CPN2 PFN2 TSC1 |
| 0.032 | gluconeogenesis | GAPDHS ENO1 |
| 0.032 | learning | APP NRXN1 |
| 0.032 | negative regulation of fibrinolysis | F2 PLG |
| 0.033 | oxidation-reduction process | QSOX1 CP TXN PRDX6 HEPHL1 |
| 0.035 | extracellular matrix organization | FGA APP SPARC COL6A3 CD44 |
| 0.040 | keratinization | KRT6B KRT9 TGM3 KRT77 KRT23 |
| 0.041 | positive regulation of transcription, DNA-templated | NRIP1 ATAD2 SKAP1 MED30 |
| 0.041 | cellular protein modification process | TGM1 TGM3 |
| 0.041 | neuromuscular process controlling balance | APP NRXN1 |
| 0.048 | post-translational protein modification | QSOX1 CP SERPINA1 FGA APP |

Note: Significant GO groups with single genes were omitted.
Abbreviations: DAS-NTT, day-adjusted scores of normo-to-tachy ratio.

**Supplemental Table 12:** Proteins at baseline of Day 2 differentiating between placebo responders (≥50% reduction in nausea) and placebo non-responders according to ROC curves.

| Genes | Protein Accessions | Protein Descriptions | *P*-value |
|---|---|---|---|
| ACTN2 | P35609 | Alpha-actinin-2 | 0.003 |
| MRPL15 | Q9P015 | 39S ribosomal protein L15. mitochondrial | 0.008 |
| GAPDHS | O14556 | Glyceraldehyde-3-phosphate dehydrogenase. testis-specific | 0.010 |
| KRT31 | Q15323 | Keratin. type I cuticular Ha1 | 0.011 |
| ACAN | P16112 | Aggrecan core protein | 0.012 |
| IGHM; | P01871;P01773 | Ig mu chain C region;Ig heavy chain V-III region BUR | 0.014 |
| TFRC | P02786 | Transferrin receptor protein 1 | 0.264 |
| NUCB1 | Q02818 | Nucleobindin-1 | 0.296 |
| FAM83G | A6ND36 | Protein FAM83G | 0.353 |
| FER1L5 | A0AVI2 | Fer-1-like protein 5 | 0.417 |
| SLC9A3R1 | O14745 | Na(+)/H(+) exchange regulatory cofactor NHE-RF1 | 0.782 |

**Supplemental Table 13:** Proteins at baseline of Day 2 differentiating between placebo responders (≥50% reduction in motion sickness) and placebo non-responders according to ROC curves.

| Genes | Protein Accessions | | *P*-value |
|---|---|---|---|
| IGKV1D-16 | P01601 | Ig kappa chain V-ID region 16 (Fragment) | 0.004 |
| IGHV3-23 | P01764 | Ig heavy chain V-III region 23 | 0.010 |
| ARHGDIB | P52566 | Rho GDP-dissociation inhibitor 2 | 0.012 |
| WDR62 | O43379 | WD repeat-containing protein 62 | 0.015 |
| MASP2 | O00187 | Mannan-binding lectin serine protease 2 | 0.017 |
| NINL | Q9Y2I6 | Ninein-like protein | 0.034 |
| | P01597 | Ig kappa chain V-I region DEE | 0.055 |
| ADCY2 | Q08462 | Adenylate cyclase type 2 | 0.072 |
| GZMB | P10144 | Granzyme B | 0.110 |
| QSOX1 | O00391 | Sulfhydryl oxidase 1 | 0.174 |
| ZNF806 | P0C7X5 | Zinc finger protein 806 | 0.227 |
| IGHG1 | P01857 | Ig gamma-1 chain C region | 0.417 |
| LRP5 | O75197 | Low-density lipoprotein receptor-related protein 5 | 0.544 |
| TXN | P10599 | Thioredoxin | 0.647 |
| MCM3AP | O60318 | Germinal-center associated nuclear protein | 0.789 |
| CP | P00450 | Ceruloplasmin | 0.824 |
| PPIP5K2 | O43314 | Inositol hexakisphosphate and diphosphoinositol-pentakisphosphate kinase 2 | 0.843 |
